# Evidence that local viscosity and NOX-dependent ROS increases render the tardigrade H. exemplaris resilient to extreme physical force

**DOI:** 10.64898/2026.05.14.724643

**Authors:** Molly J. Kirk, Jonathan Paules, Samantha L. Fiallo, Amelia M. Leeman, Carl D. Meinhart, Joel H. Rothman

## Abstract

Biological phase changes provoked by stress, such as vitrification or gel-sol transitions, enable many organisms, including extremotolerant tardigrades, to enter quiescent states and survive extreme environmental conditions. Protein-driven phase transitions are hypothesized to produce large-scale changes in intracellular viscosity, allowing tardigrades to survive extreme stresses such as desiccation. We report that the tardigrade *Hypsibius exemplaris* undergoes both large-scale and local increases in intracellular viscosity following exposure to anoxic and hyperosmotic stress. Such dramatic shifts in cellular viscosity would be expected to enhance cellular resilience to physical force. Indeed, we found that tardigrades can survive, behave normally, and reproduce after exposure to the highest simulated hypergravity (HG) achievable in an ultracentrifuge (one million times Earth’s gravity). In contrast, *Caenorhabditis elegans*, a similarly sized animal, does not survive these extreme forces owing to loss of cellular integrity. Remarkably, tardigrades frozen during exposure to extreme hypergravitational force show minimal disruption of fine cellular ultrastructure and little evidence of stratification of cellular components whose density varies by nearly a factor of two. Further, exposure to anoxia, hyperosmotic stress, and HG all result in a large increase in reactive oxygen species (ROS), which is required for survival under these extreme environments. Inhibition of NADPH oxidase (NOX) suppresses survival both to HG and hyperosmotic stress. Our findings suggest that intracellular viscosity changes in response to multiple extreme stresses may underlie the resilience of these animals to extraordinary physical stress, and that survival in or recovery from these states relies on ROS signaling via NADPH oxidase.

**Significance Statement:** Tardigrades are renowned for surviving conditions that are lethal to nearly all other life forms. We reveal two mechanisms that support this resilience: intracellular viscosity changes and NADPH oxidase-mediated ROS signaling. Through direct assessment of the effects of altered cellular material properties, found that tardigrades are resilient to forces up to one million times Earth’s gravity, establishing them as the most hypergravity-resistant animal currently known.

## Introduction

Extremotolerance, or the capacity of an organism to withstand extreme environmental conditions, has convergently evolved across many phyla in response to a broad range of externally imposed stresses (1–4). Within the animal kingdom, a foremost example of extremotolerance is seen in the phylum Tardigrada, animals renowned for their resistance to extraordinarily high ionizing radiation (5), freezing (6), near-complete desiccation (7), anoxia (8), and even the vacuum of space (9). This breadth of resilience to extremes establishes tardigrades as foundational models for investigating the environmental limits of animal life.

While the tardigrade’s physiological response differs across various stressors, some common themes have emerged. Transcriptomic analysis of the desiccated (anhydrobiotic, or “tun”) state of the tardigrades *Hypsibius exemplaris* and *Ramazottious varieornatus* revealed that production of reactive oxygen species (ROS) scavenging enzymes is a key biological program induced by anhydrobiosis (10, 11). Further, transition into the desiccated state results in increased ROS production. As a result of this burst of these damaging molecular species, ROS-scavenging enzymes are required for survival under desiccation stress in the tardigrade *Paramacrobiotus spatialis* (12). However, in *H. exemplaris*, ROS production is also required for entry into the osmotic stress-induced tun, as quenching of ROS prevents tardigrades from entering the tun state, and treatments that increase ROS production promote tun formation in the absence of osmotic stress (13). Thus, ROS production may serve as a key signaling system for the induction of the stress response, but ROS-scavenging enzymes are required to maintain homeostasis and prevent significant oxidative cellular damage. This dual nature of ROS as a signal for stress responses and an inducer widespread cellular damage if not appropriately regulated, has been observed across biology, including in other extremobionts (2), cancer, and bacterial infection (14, 15).

Another common feature that appears to underlie stress resilience in tardigrades is the use of phase transitions (16, 17), which are hypothesized to support biological structures under extreme stress (18). In contrast to many other extremotolerant organisms, which increase intracellular viscosity through accumulation of disaccharides and subsequent vitrification leading to biostasis, tardigrades produce minimal levels of vitrifying disaccharides (19). Instead, these animals have evolved tardigrade-specific intrinsically disordered proteins (Tardigrade Disordered Proteins or TDPs) that undergo biophysical phase transitions, forming fibrous or punctate structures. These structures have been observed both *in vitro* and when expressed in human cells (18, 20, 21). Different classes of TDPs localize to specific cellular and extracellular compartments (22), where they are hypothesized to support cytoplasmic and organellar structures in the absence of water (23), potentially playing a role in metabolic suppression through large-scale alterations in molecular diffusion and flow rates (18). Such phase transitions would be expected to cause large-scale changes in molecular motion by altering macro-viscosity, the bulk state of a cell, and micro-viscosity, experienced at the level of individual small molecules (24). However, changes in intracellular viscosity and how such phase transitions might alter material properties and physical resilience of tardigrade cells have yet to be investigated in response to any extreme environmental stressor.

Here we report that in response to anoxia and hyperosmotic stress, tardigrades experience up to a 15-fold increase in intracellular cytoplasmic viscosity. Consistent with their capacity to undergo such large-scale increases in viscosity, we found that active-state tardigrades can survive relative centrifugal forces equivalent to 1 million times the force of Earth’s gravity (1M x *g*), conditions that are sufficient to sediment large macromolecular complexes in living cells (25). Remarkably, the fine ultrastructure of animals frozen under such extreme hypergravity (HG) shows no salient evidence for density-dependent fractionation of cellular components, despite the ∼2-fold difference in component density within cells (26, 27). These findings suggest that the dramatic increase in intracellular viscosity may promote resilience to extreme physical force and density-dependent fractionation. Further, we found that animals recovering from this extreme HG showed a significant increase in ROS levels and that inhibition of ROS production prevented the successful transition into, survival in, or recovery from HG, anoxia, and hyperosmotic stress. Pharmacological inhibition studies implicate NADPH oxidase in this ROS production and subsequent survival in osmotic and HG stress response. Collectively, these findings suggest that the increases in cellular viscosity occurring in response to multiple stresses prevent density-dependent fractionation at 1 million x g, and that these stress responses require ROS signaling to promote survival to such extreme physical forces.

## Results

### Exposure of tardigrades to extreme environments leads to a precipitous increase in large-scale intracellular viscosity

While it has been hypothesized to occur, direct measurement of *in vivo* viscosity changes induced by extreme stressors has not yet been reported in tardigrades. We therefore sought to investigate the occurrence and magnitude of intracellular viscosity changes in *H. exemplaris* under conditions of osmotic and anoxic stress (Fig. 1a). We first assessed survival rates under exposure to these stressors as a function of time and found that the animals can survive >16 hours of osmotic stress (0.5 M D-mannitol, pH 7.0) (Fig. S1a) and >24 hours in response to either N_2_- or CO_2_-promoted anoxia (0.05 +/- 0.028 ppm D.O., pH 7.0 or 0.045 +/- 0.006 ppm D.O., pH 3.4, respectively) (Fig. S1b and c), as summarized in Fig. 1b. Our subsequent analyses were performed on quiescent animals over time courses in which >75% of animals can reanimate and fully recover motor function.

**Figure 1:**
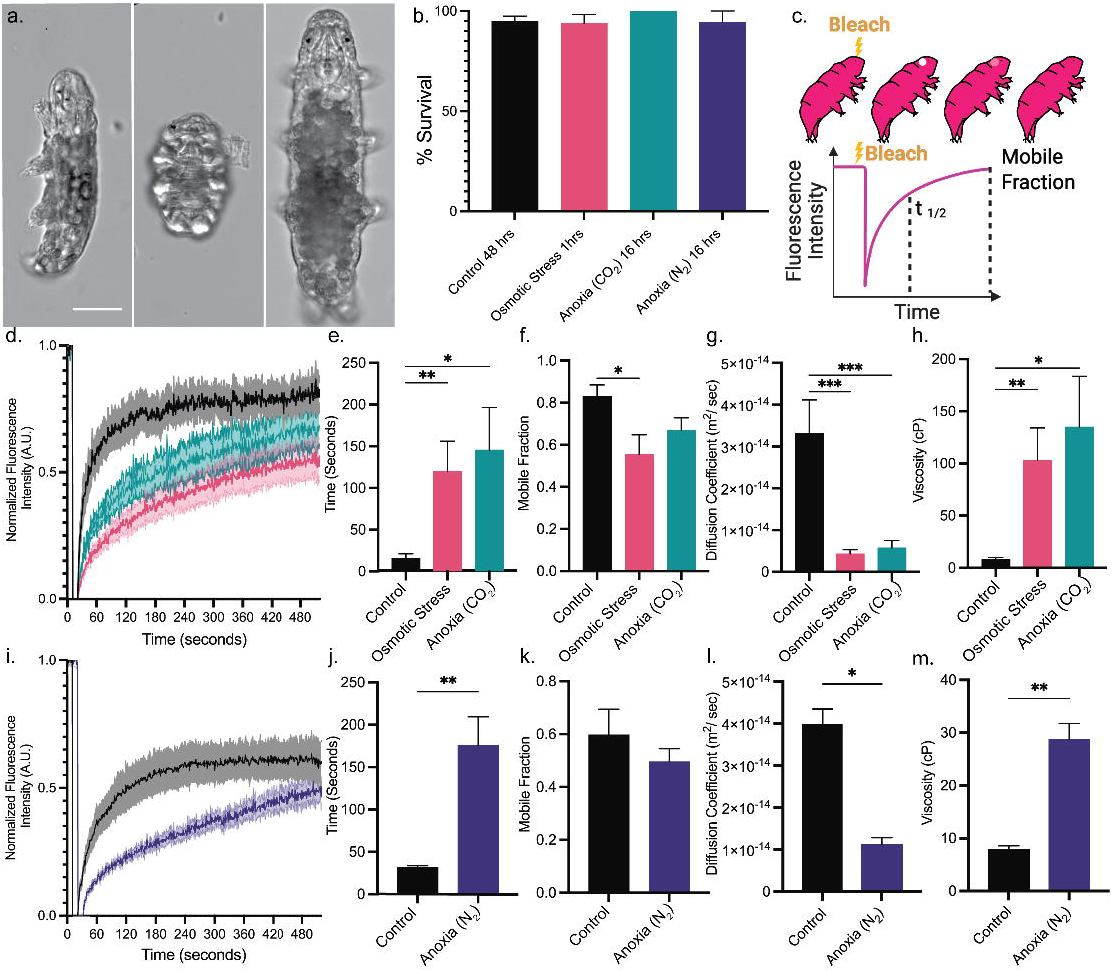
Tardigrade intracellular viscosity is modulated by exposure to extreme environments. a) DIC microscopy images of tardigrades under rearing conditions (left), 0.5 M mannitol (center), and anoxic conditions (right). Scale is 50 μm. b) Graph of percent survival rates for animals under extreme stressors, shown above, is the maximum time points where over 75% of the animals survived. c) Schema of derived metrics from FRAP experimentation. d) Raw averaged traces, with S.E.M. error inlay, of Fluorescence Recovery taken from animals under rearing conditions, osmotic, and anoxic stress. e) Measured tau at half maximum values in seconds for fluorescence recovery after photobleaching from a 2.5 μm circular bleach in the head region, **, p=, 0.0078 *, p=0.0414. f) Mobile fraction of dye under rearing conditions, osmotic stress, and anoxia *, p=0.0305 g) Coefficient of Diffusion calculated using Soumpasis model ***, p = 0.0009 and 0.0021. h) Macro-viscosity calculated in centi-poise (cP) using the Einstein-Stokes equation for a tumbling ellipse in 3 dimensions **, p=0.0039 and *, p=0.0196. i) Raw averaged traces, with S.E.M. error inlay, of Fluorescence Recovery taken from animals under normoxic 20mM Lactate buffer and anoxic conditions. j) Measured tau at half maximum values in seconds for fluorescence recovery after photobleaching from an 8 μm x 5 μm rectangular bleach in the head region, **, p=0.0036. k) Mobile dye fraction under normoxic 20mM Lactate buffer or anoxic 20 mM Lactate buffer, no significant difference. l) Coefficient of Diffusion derived using SIM-FRAP, *, p = 0.0159. m. Macro-viscosity calculated in centi-poise (cP) using the Einstein-Stokes equation for a tumbling ellipse in 3 dimensions *, p=0.0336. All data presented as mean with S.E.M.

To evaluate large-scale intracellular viscosity in these quiescent states, we monitored cytoplasmic mobility of mitochondria labeled with MitoTracker Deep Red FM by Fluorescence Recovery After Photobleaching (FRAP) (28)(Fig. 1c). Osmotic stress induces a rapid (<10 min) and pronounced morphological transition into a tun, during which the animal contracts along the anteroposterior axis, the body volume shrinks, and the limbs retract into the body. Accompanying this dramatic transformation, the animals become quiescent, with no sign of physical activity. During exposure of *H. exemplaris* to osmotic stress, we observed a large increase in fluorescence recovery time over a 5 min time-course following laser-bleaching of a 2.5 μm diameter circle in the head region (see Materials and Methods). This response resulted in a 7.6-fold increase in time to half-maximal recovery (τ_1/2_) compared to non-stressed controls (Fig. 1d, e; p=0.0078). To further evaluate this decrease in apparent mobility of mitochondria, we monitored the mobile fraction, a measure of the proportion of mitochondria that are free to diffuse into the bleached region during the recovery period (see Materials and Methods for equations and derivations) and observed a significant decrease (∼23%; p=0.0387) in the mobile fraction under osmotic stress (Fig. 1f). Using the FRAP measurements, we applied a Soumpasis model (see Materials and Methods) to derive a diffusion coefficient and found that it was substantially diminished (87% lower, p = 0.0009, Fig. 1g), than in control animals, indicating reduced diffusion rates in the osmotically stressed tun state, consistent with the animal’s quiescence.

As with osmotic stress, exposure to anoxia also induces rapid quiescence in tardigrades. However, oxygen deprivation results in a pronounced difference in morphology: the animal undergoes elongation over the course of 10 mins, and shows a dramatic increase in volume, with the limbs extending rather than retracting. Despite the contrasting response of oxygen starvation compared to that of osmotic stress, exposure of the animals to CO_2_-sparged water resulted in a large increase (9.3-fold, p=0.0414) in the τ_1/2_ FRAP recovery (Fig. 1 d and e). In contrast to osmotic stress, anoxia (CO_2_) does not significantly change the mobile fraction compared to that in non-stressed, active animals (Fig. 1f).

To extend these findings to a second more physiologically relevant anoxic stress, we sparged a serum vial containing 20 mM sodium lactate buffer with ultrapure nitrogen prior to the addition of Oxyrase (29), an enzyme that selectively converts molecular oxygen to water. By this method, dissolved oxygen levels were reduced to 0.05 PPM +/- 0.028 S.E., similar to that seen in CO_2_-sparged water (Fig. S2). Under these conditions, we observed that tardigrades immediately cease movement and, over the course of 10 mins, achieve a similar morphology to that of CO_2_-exposed animals (Fig. 1a). The anoxic vial increased imaging difficulty requiring 10x magnification and bleaching of a rectangular area (8 μm x 5 μm) to assure sufficient bleach depth (Fig. 1i). Under these conditions, the FRAP τ_1/2_ was increased 6-fold compared to that in the normoxic lactate buffer (Fig. 1j). However, in contrast to what we observed for osmotic stress, we found that the mobile fraction of mitochondria in both anoxic conditions were statistically similar (p=0.3189) to that under normoxic conditions (Fig. 1k).

These observations indicate that osmotic stress both generally restricts the mobility of mitochondria and results in sequestration/immobilization of a subpopulation of the organelles, resulting in a decrease in mobile fraction. In contrast, while anoxia also leads to a general decrease in mitochondrial mobility (increased τ_1/2_), it does so without significantly sequestering a subset of mitochondria (i.e., no change in mobile fraction; Fig. 1f and j).

To evaluate the magnitude of viscosity changes from these experiments, we applied the Einstein-Stokes Equation to estimate the apparent viscosity experienced by a mitochondrion-size structure of 0.5 μm x 0.5 μm x 1 μm, modeled as a three-dimensional tumbling ellipse (30) for circular bleach patterns and used the SIM Frap FIJI plugin for rectangular bleach patterns (see Materials and Methods). These estimations revealed a dramatic change in viscosity under the stressed conditions. Under hyperosmotic stress, we observed a 12-fold increase in apparent viscosity (average of 103 cP for experimental compared to 8 cP for control; p=0.0039). Anoxic stress induced by CO_2_ resulted in 15-fold increase in apparent viscosity (average of 125 cP for experimental; p = 0.0196; (Fig. 1h). Finally, induction of anoxia with N_2_ resulted in a 3.6-fold increase (average of 28.6 cP compared to 7.8 cp in controls; Fig. 1m) (Diffusion coefficients for these samples are shown in Fig. 1l). It should be noted that induction of anoxia CO_2_ reduced Mitotracker Deep Red fluorescence intensity *in vivo* (Fig. S3), suggesting that mitochondrial membrane potential may be affected by anoxic CO_2_ conditions, despite the dye’s relative insensitivity to this parameter (31). Nevertheless, the diffusion kinetics of dye-labeled mitochondria is independent of dye intensity, owing to within-sample normalization (32).

We conclude that both extreme environments, osmotic stress and anoxia, greatly increase intracellular viscosity, however the fraction of mobile mitochondria is significantly lower in the latter state, suggesting enhanced caging under this condition.

### Micro-scale increase in viscosity induced by extreme stress

While our findings demonstrate that viscosity increases sharply *in vivo* in response to two unrelated extreme stresses, they do not reveal the spatial scale of this effect. If the observed viscosity increase is sufficient to drive a loss of metabolic flux, it would be expected to alter micro-viscosity, reflecting resistance to flow at the level of single molecules (Fig. 2a). To assess possible changes in micro-viscosity, we used the molecular rotor dye Viscous Aqua (VA, URSA Biosciences), which functions essentially as a molecular motion sensor. At low local viscosity, two parts of the molecule freely rotate around a single bond, preventing π-bond conjugation that is required for a fluorescent signal. However, as the micro-viscosity increases, the dye becomes stabilized in the planar form, thereby increasing its fluorescence intensity inversely to the slowing of molecular motion (33) (Fig. 2b).

**Figure 2:**
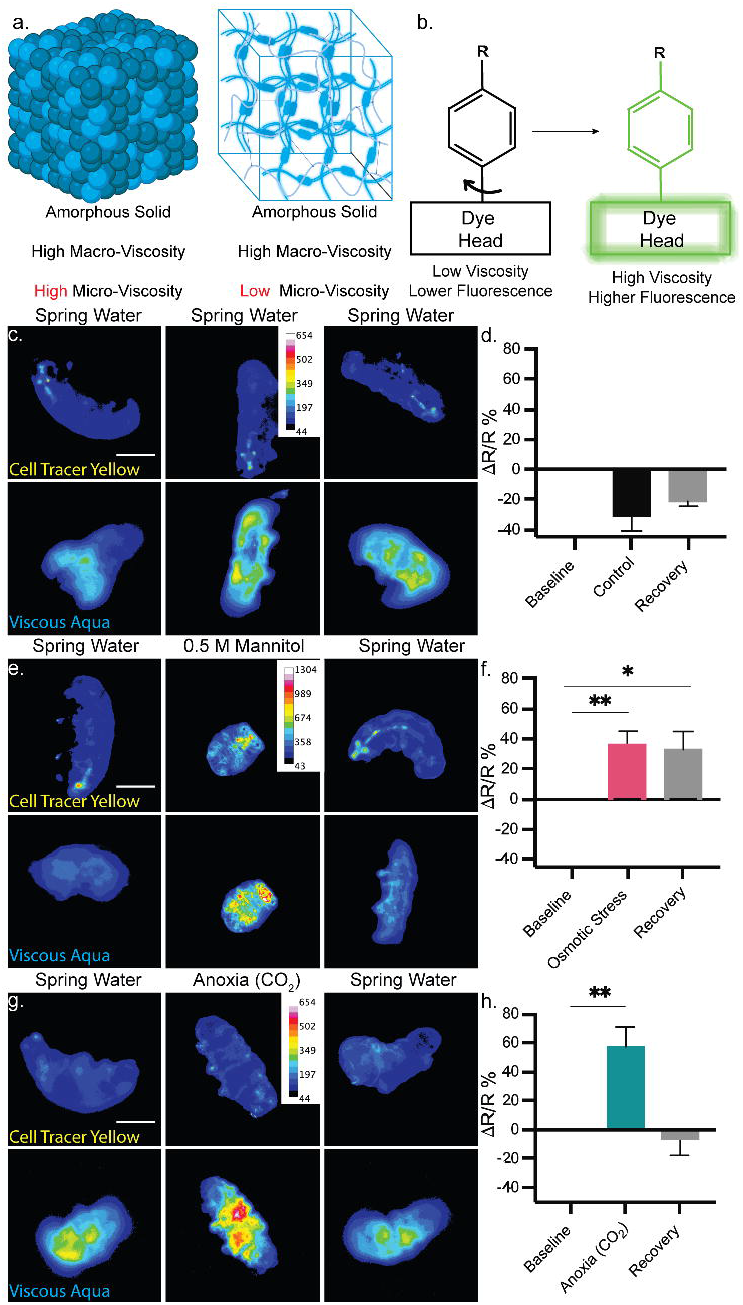
Intracellular local viscosity is modulated by exposure to extreme environments. a) Schema showing two proposed models for amorphous solid formations caused by TDPs in response to anhydrobiosis, one with molecular crowding and one without. b) Schematic description of molecular rotor dye function. c) Spring water control micrographic heat maps of CTY and VA fluorescence intensity, before, during, and after exposure to inert spring water. d) Quantification of %Δ*R*/*R* where R is the VA fluorescence intensity over CTY fluorescence intensity. All comparisons were non-significant. e) Intensity heatmap micrograph of the animal before, during, and after exposure to 0.5 M Mannitol in spring water. The scale for all images is 50 μm. f) %Δ*R*/*R* quantification of mannitol exposed animals, *, p=0.0233 and **, p=0.0014. g) Intensity heat map of tardigrades before, during, and after 10-minute exposure to 0.5 ppm +/- 0.1 CO_2_ sparged water, pH 3.4. h) Percent change in fluorescence ratio, of VA and CTY, before, during, and after exposure to CO_2_ sparged water, **, p = 0.0016. All data presented as Mean and S.E.M.

To assess changes in micro-viscosity, we imaged tardigrades stained with VA before, during (10 minutes after onset of stress), and after (10 minutes after recovery) exposure to stress conditions and compared the signal to that in control animals suspended in spring water alone (Fig. 2c). We controlled for volumetric changes that could alter fluorescence intensity measurements using Cell Tracer Yellow (CTY), a cytosolic dye that is not viscosity sensitive. Micro-viscosity changes were monitored by analyzing the VA:CTY ratiometric fluorescent signal which, under control conditions, showed no significant change (Fig. 2d).

These analyses revealed a statistically significant and persistent increase (average 36%; p=0.0014) in fluorescence ratio under osmotic stress (Fig. 2e and f). Following a return to normal osmotic conditions and a 10-minute recovery period, the VA:CTY ratio did not return to baseline (average increase of 33%; p=0.0233), implying that the increased micro-viscosity remained high following recovery. This observation is consistent with significantly decreased motor output of the animals, as compared to controls.

We similarly examined changes in micro-viscosity during anoxia by immersing the animals in CO_2_-sparged water (Fig. 2g). In this environment, we observed a sharp increase in the VA:CTY ratio (average 58%; p=0.0016). However, in contrast to our observations with osmotic stress, we found that the VA:CTY ratio was restored to baseline levels following a ten-minute recovery period (-7.063%; p=0.546; Fig. 2h), consistent with the return of the animals to normal activity levels and rate. The increase in VA:CTY ratio suggests either a substantial increase in intracellular micro-viscosity or an artifact in VA fluorescence resulting from removal of triplet state quenching under low oxygen conditions (34). To investigate this possibility, we performed spectroscopy of VA under anoxic conditions and observed no significant change in VA normalized fluorescence intensity in CO_2_-sparged water (1.07 fold control intensity, p=0.9704), or in Oxyrase-treated anoxic lactate buffer, which had a dissolved oxygen content of 0.17 +/- 0.01 ppm S.E. (0.96 fold control intensity, p=0.9993; see Materials and Methods) (Fig. S4). This finding indicates that the anoxia-induced increase in VA fluorescence is not attributable to suppression of quenching by oxygen. While it was conceivable that the lower pH (3.4) of the CO_2_-sparged water alters the fluorescent behavior of VA, we found that exposure of dyed animals to a pH 3 sodium citrate buffer did not result in significant changes in the VA:CTY ratio *in vivo* (Fig. S5).

These findings suggest that both osmotic and hypoxic stress induce rapid, precipitous increases in micro-viscosity within 10 mins of exposure. However, the kinetics of the return to baseline micro-viscosity are state-dependent, with osmotically stressed animals maintaining high viscosity even 10 mins into recovery. Regardless, the sharp increase in macro- and micro-viscosity in the animals implies that resistance to bulk flow is greatly elevated and intracellular molecular motion reduced in response to both types of stress.

### Tardigrades are resilient to extreme centrifugal forces of up to 1,000,000 x g

Having observed osmotic and hypoxic stress-induced increases in both macro- and micro-viscosity, we sought to test whether these changes in *in vivo* material properties might enhance cellular and organismal resilience to extreme physical forces. Given that elevated viscosity should reduce macromolecular and organellar movement within cells, we posited that tardigrades may be resilient to the extreme physical force imposed by simulated hypergravity (HG).

In most multicellular organisms, sustained HG generally leads to rapid lethality. 9 x g is lethal to humans within minutes, even when protected by G-suits and performing anti-G-straining maneuvers during flight, with consciousness typically lost at ∼ 5 x g (35). As HG resilience is inversely related to animal size, diminutive animals can survive much higher levels of HG. However, 9,000 x g is lethal to adult *Drosophila melanogaster*, among the most g-force resilient arthropods (36). Small extremotolerant animals have been shown to be more resilient to HG. For example, *C. elegans* can survive 400,000 x g for 45 minutes, after which no visible changes to morphology or reproduction were observed (37). Moreover, the tardigrade *H. exemplaris* has been reported to be survive at least 16,060 x g before death of any animal was seen (38). Importantly, owing to the lack of existing technology at the time of publication, neither the *C. elegans* nor *H. exemplaris* studies assessed the maximum g-force that can be sustained by either animal (37, 38). An overview of maximum HG resilience across the tree of life can be found in Supplemental Table 1.

To investigate the limits of resilience of tardigrades to such extreme physical stress, we analyzed the survival of active-state *H. exemplaris* when subjected to various g forces for 10 mins in an ultracentrifuge (Fig. 3a; see Materials and Methods). Remarkably, virtually all animals survived HG of up to 675,000 x g, the greatest force that any metazoan has been reported to withstand (39). Nearly all these animals recovered and showed largely normal behavior following HG. The maximum HG currently attainable in a modern ultracentrifuge is ∼1,000,000 x g (1M x g) (40). We found that ∼50% of active-state animals survived when exposed to 1M x g for 10 mins (Fig. 3b, p=0.0437). This observation is particularly noteworthy, as this g-force is sufficient to sediment ribosomes and glycogen within living vertebrate cells, ultimately resulting in near complete stratification and cell lysis within minutes (25).

**Figure 3:**
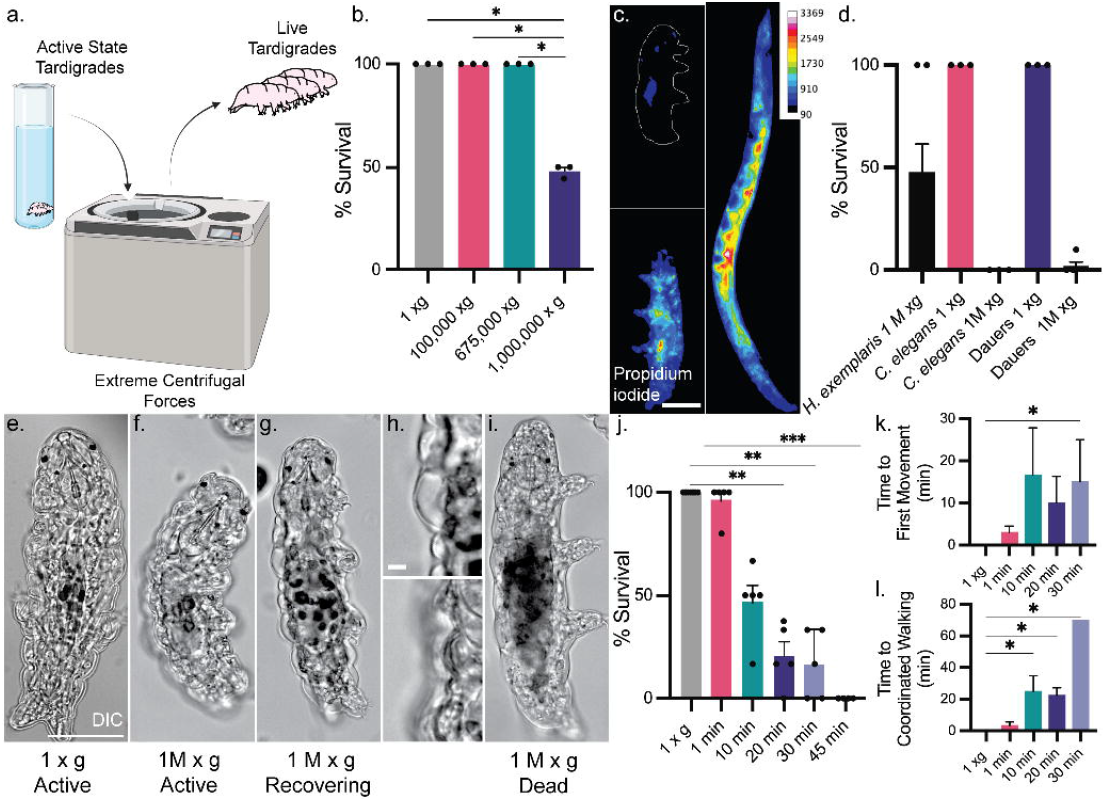
Tardigrades are resilient to 1M x g for over 30 minutes. a) Schematic of experimental design. b) Survival rate of tardigrades exposed to 1 x g, 100,000 x g, 675,000 x g, and 1,000,000 x g for 10 minutes at maximum g-force. All survival data represent the survival rate among six animals per trial run across at least 3 experimental days, *, p = 0.0437. c) Intensity heat map of surviving Tardigrade (left, top), non-surviving tardigrade (left, bottom), or adult *C. elegans (right)* centrifuged in the presence of propidium iodide. Scale is 50 μm. d) Survival rate of tardigrades, and adult and dauer *C. elegans* following 10-minute centrifugation at 1M x g, *, p=0.0323. DIC micrograph of active state (e.), active state post centrifugation (f.), recovery state (g.), zoomed in inlay of recovery state (top, h) and recovered state (bottom, h), and post centrifugation dead animals imaged 20 minutes after return to 1 x g (i.). The scale is 50 μm. j) Percent survival following 1M x g exposure for 1, 10, 20, 30, and 45 minutes, **, p=0.0035 and 0.0016, and ***, p=0.0003. Average of time to first motion (k.), *, p=0.035, and time required to reach coordinated motion (l.) as a function of length of HG exposure, *, p = 0.0194, 0.0181, and 0.0175, respectively. All data presented as mean with S.E.M.

To test whether the extreme HG results in cell lysis or loss of cellular integrity, we repeated the centrifugation in the presence of propidium iodide, a cell death indicator. We first evaluated the efficacy of propidium iodide in detecting dead cells and loss of membrane integrity within the animals by subjecting them to a lethal high temperature without centrifugation (50°C for 45 minutes, Fig. S6) and found that heat-killed animals filled with the dye, confirming its ability to detect cells that have lost integrity. After centrifugation, non-surviving animals showed loss of cellular integrity (Fig. 3c, left bottom), while in surviving tardigrades few cells stained positive for the dye (Fig. 3c, left top). These findings indicate that plasma membrane integrity is maintained during and following HG in surviving animal populations (Fig. 3d).

The ability of an animal with thousands of cells to withstand 1 M x g prompted us to consider whether another diminutive animal, the nematode *Caenorhabditis elegans*, which is similar in mass and cell count to tardigrades, and which is reported to resist 400,000 x g (37), could also endure 1 M x g. However, when exposed to 1M x g for 10 minutes, 100% (n = 50) of day 1 adult *C. elegans* died (Fig. 3d) as indicated by their elongated body plan and lack of detectable motion over the course of 24 hours. Following centrifugation in the presence of propidium iodide, all adult *C. elegans* cells showed strong staining, in contrast to the lack of staining observed in tardigrades (Fig. 3c). These observations suggest that the ability to survive extreme HG is not a characteristic of an animal of similar size and cell composition to that of a tardigrade but may be a distinct characteristic of tardigrade physiology.

An alternative developmental stage of *C. elegans* that is induced by, and particularly resilient to, several stresses, including desiccation and extended periods of starvation, is the dauer larva (41). We found that while this more resilient form of the animal initially survived following centrifugation at 1 M x g, evident from their mobility, animals were no longer moving and assumed an elongated body plan after approximately one hour. Only one animal out of 50 exposed to HG recovered from the dauer state 24 hours after centrifugation, showing normal return to the L4 stage (Fig. 3d) and progressed to adulthood and produced viable progeny. These observations reveal that the specialized *C. elegans* dauer state may be resilient to HG exposure owing to the sealed cuticle structure and resistance to other stressors such as desiccation (41); however, it appears less so than are active tardigrades.

Following exposure of tardigrades to extreme HG, we observed three distinct classes of behavior and morphology. In the first class, some animals showed coordinated, albeit slower, locomotion immediately after removal from the ultracentrifuge (Fig. 3f), with minimal observable differences from the 1 x *g* controls (Fig. 3e). In the second class, animals did not show movement upon return to 1 x *g* (Fig. 3g). In these animals, the epidermis initially appeared retracted from the cuticle and the overall morphology resembled the elongated form seen during anoxia and immediately following recovery from hyperosmotic stress (Fig. 3h, top). Animals in this class recovered with restored coordinated motion within 20-30 minutes of return to 1 x *g* (Fig. 3h, bottom). The third class initially appeared identical in morphology to the second class, but the animals never recovered locomotor function, remaining in an extended morphology (Fig. 3i) and ultimately dying.

We assessed the impact of time of exposure to 1 M x *g* on survival rate. Remarkably, we found that while lethality increased with longer exposures, some tardigrades survive 1 M x *g* even after 30 minutes of HG. At longer times under HG, the survival rate dropped to zero (Fig. 3j). These results suggested an exponential loss of survival with exposure time to HG (Fig. S7). Further, time to recovery positively correlated with exposure duration based both on time to first detectable movement (Fig. 3k) and time to coordinated walking motion (Fig. 3l). Finally, we investigated the long-term impact of extreme HG by monitoring survival rate (Fig. S8a), egg production (Fig. S8b), and progeny number (Fig. S8c) of the animals over the course of the subsequent 16 days. While HG-exposed survivors showed shorter lifespans, they nonetheless reproduced successfully, even after 30-minute exposure to 1M x *g*. Our results indicate that active-state tardigrades are resilient to extreme HG forces over an extended period, recover in an exposure-dependent manner, and retain reproductive capacity following HG exposure, a level of resilience to extreme physical force that has not been previously reported for any animal.

### HG-induced viscosity changes may result from effects of a coincident stress response

As extreme HG does not occur naturally on Earth, tardigrades would not have evolved specific mechanisms to withstand such forces. Rather, it is likely that this phenomenon results from their ability to switch into states that are resilient to extreme environments encountered in nature. To evaluate this possibility, we assessed whether other stresses enhanced survival to HG. We found that anoxic stress did not substantially enhance the ability of the animals to withstand HG: when subjected to ultracentrifugation at 0.045 +/- 0.01 ppm atmospheric oxygen (see Materials and Methods), we observed a minor (10%), and non-significant, increase in survival rate compared with normoxic conditions (Fig. S9a).

In contrast, however, we found that pre-exposure to high osmotic conditions, which induces the tun state, greatly increased survival rates at 1 million x g (Fig. S9a), extending survival to over 2 hours (Fig. S9b). These findings are consistent with our observations that hyperosmotic stress substantially decreases organellar mobile fraction, while anoxia induces a state that may be more akin to a highly viscous liquid, restricting flow rate but not the fraction of mobile organelles. It is therefore conceivable that HG exposure in the active state may mimic the viscosity changes occurring in hyperosmotic stress.

To assess this hypothesis, we imaged VA-stained active-state animals following HG exposure. We observed a 43% increase in ΔR/R, indicating an increase intracellular microviscosity (Fig. 4a and b), which was maintained for at least 10 minutes after return to 1 x g (Fig. 4c). The similar magnitude and kinetics suggest that HG induces a viscosity increase that may be more similar to that of osmotic stress than anoxia. However, specimens frozen during HG (see below) showed no net alteration to overall animal volume, suggesting that overall water loss may not be the trigger for this viscosity change (Fig. S10).

**Figure 4:**
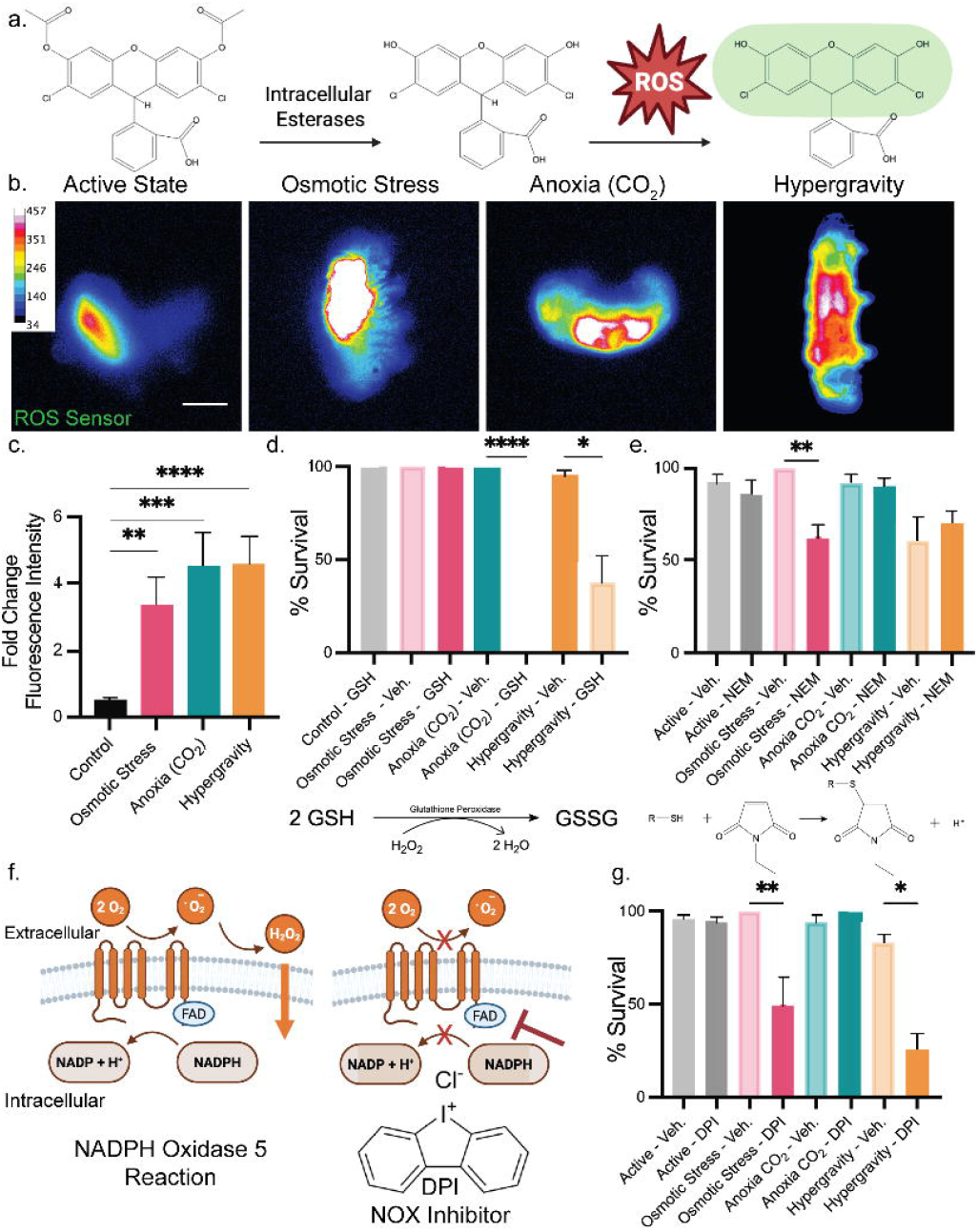
Tardigrades show viscosity change in response to HG and little to no ultrastructural density-dependent fractionation. a) Schematic of experimental outputs for exploration of anatomy following HG exposure. b) Normalized intensity heat map of fluorescence intensity before and after centrifugation. c) % change in R across a mixed population before and after exposure to 1M x g or a 1 x g control. d) Dorsal aspect (top) and mid frontal plane (bottom) DIC micrograph of an active state animal having survived HG exposure at 675k x g. e) Dorsal aspect (top), a mid-frontal plane (bottom) DIC micrograph of tardigrades that did not survive centrifugation. The scale is 50 μm. f) Frequency histogram of storage cell location in counts per quartile as a function of the dorsal-ventral axis of the animal,****, p >0.0001. Data represents mean with S.D. g) Quantification of mean cell location in quartiles, Mann-Whitney, *, p =0.0159. Data represents mean with S.D. h) TEM micrograph of gross animal ultrastructure from control (1 x g, top) animals, unperturbed regions of HG exposed animals (675k x g, middle), and perturbed regions of HG exposed animals (675k x g, bottom). Scale is 5μm. i) Representative micrographs of cuticular structure from control (top) animals, unperturbed regions of HG-exposed animals (middle), and retracted cuticles of HG-exposed animals (bottom). Scale is 2 μm. j) Representative electron micrographs of nuclear compartments, from control (top) animals, unaffected nuclei of HG-exposed animals (middle), and altered nuclei from HG-exposed animals (bottom). Scale is 1 μm. k) Micrographs of representative mitochondrial compartments revealing normal mitochondrial cristae structure at 1x g (top), a blank panel showing no unperturbed mitochondria were visualized in this data set (middle), and a representative TEM micrograph of mitochondria observed in animals exposed to HG (bottom). Scale is 0.5 μm. l) Quantification of data represented in panel (i.), showing number of retracted cuticle events across all specimens imaged. 1 x g N=10, 675k x g N=9). m. Quantification of nuclear deformation frequency, showing total counts of nuclei visualized per condition, total number of abnormal nuclei visualized per condition, percent of deformed nuclei per condition, and sample number representing individual animal samples per condition. Data represents mean with S.E.M.

### Exposure to extreme HG causes minimal disruption of cellular ultrastructure and no evidence for density-dependent fractionation

A range of eukaryotic cells have been shown to undergo density-dependent fractionation when exposed to centrifugal forces (reviewed in (25)). As would be expected, the low-density components migrate toward the centripetal side of the cell and dense structures toward the centrifugal (radially outward) direction, resulting in stratification of these cellular components (25). Structures that are denser than the general internal cellular milieu (1.05-1.1 g / ml; (27)), such as the nucleolus (∼1.7-2.0 g/ml; (26)), and those that are less dense than the cytoplasm, including lipid droplets (∼0.9-1.0 g/ml; (27)) would be expected to move in opposite directions in the HG field, as we modeled using a COMSOL Newtonian cell model (Fig. S11). Forces exceeding 150 000 x *g* are sufficient to bring about stratification and cell rupture in even the smallest eukaryotic cells, which are 5-7 µm in diameter and similar in size to tardigrade cells (25). (Further description of density-dependent fractionation of cells and within organelles *in vivo* and *in vitro* can be found in Supplementary Table 2 and Supplementary Table 3.)

As an initial assessment of potential density-dependent stratification in tardigrades exposed to extreme centrifugal forces, we first monitored localization of the free-floating, lipid-rich storage cells by DIC optics. Following a 30-minute centrifugation at 675,000 x g, 78% of the animals survived, as evident from their mobility after return to 1 x *g* (Fig. S12); in these animals, the storage cells were distributed normally across the dorsal-ventral axis, as assessed by quantifying their abundance in four quadrants (Fig. 4 d, e, f). In animals with no sign of movement 20 minutes after centrifugation, the distribution of the storage cells was heavily biased toward the dorsal aspect of the animal (Fig. 4g), suggesting that these lighter cells were displaced upward in the centrifugal field in the non-surviving animals. This shift in the position of storage cells does not reflect their differential loss or fusion, as their number and relative size were the same between survivors and non-survivors following centrifugation (Fig. S13).

To assess the extent of density-dependent stratification at the cellular ultrastructural level, we performed Transmission Electron Microscopy (TEM) on tardigrades that were frozen during exposure to HG and fixed immediately after return to normal gravity (see Materials and Methods). Comparison of specimens subjected to HG animals revealed that cellular ultrastructure was largely unaltered by this extreme physical stress, remaining largely similar to controls. We observed little evidence of density-dependent stratification of cellular structures. At the gross anatomical level, the only visible stratification we observed was dislocation of electron-dense vacuoles of reserve material, a known identifier of type I storage cells, (42) indicating partial intracellular stratification of storage cell in these animals (Fig. 4h, bottom).

While TEM revealed that the overall cellular architecture was largely unchanged, we did observe some disruption of structure following exposure to extreme HG. Specifically, we found a greater frequency of cuticular-epidermal separation in animals exposed to HG compared to those frozen at 1 x *g* (Fig. 4i, quantified in panel 4l, a 16.67-fold increase in separation frequency, p=0.0341). Further, in contrast to control animals, in which all nuclei showed a normal morphology, the nuclei of centrifuged animals appeared elongated in rare cases (∼3%, 13 nuclei of the 415 nuclei examined), and the nucleolar compartment formed an ellipsoid. In these nuclei, the surrounding nucleoplasm was less electron-dense, and strands were observed to radiate from the nucleolus (Fig. 4j, quantified in Fig. 4m). Mitochondria showed the most pronounced and consistent ultrastructural perturbation, with poorly defined cristae in all cells examined (Fig. 4k).

Collectively, these results indicate that the ultrastructure of tardigrades exposed to extreme centrifugal forces remains grossly unperturbed, with scant evidence for stratification of cellular components and minimal changes to cuticular and nuclear morphology. However, the perturbed mitochondrial morphology suggests potential alteration of mitochondrial function.

### Evidence that reactive oxygen species (ROS) originating from NADPH oxidase are required for survival in hypergravity and hyperosmotic stress

Reactive oxygen species (ROS) can be produced from both mitochondrial and cytosolic sources and serve various roles within cells (43). Recent studies reveal that ROS signaling mediates survival under desiccation and hyperosmotic stress in tardigrades (12, 13). However, the role of ROS production in response to low oxygen or HG has not yet been assessed. Given the significant disruption of mitochondrial ultrastructure during exposure to HG, we first sought to assess whether ROS is induced in response to HG. Indeed, we observed a 4.5-fold (p < 0.0001) increase in fluorescence intensity of the ROS-sensitive dye 2′,7′-dichlorodihydrofluorescein diacetate (DCF-DA) within animals following 10-minute exposure and 10-minute recovery from 1 M x *g* HG (Fig. 5a, b, and c). We note that, as oxidation of DCF-DA by ROS is an irreversible reaction, the rate of ROS clearance is not revealed in this experiment (Fig. S14, (44)).

**Figure 5:**
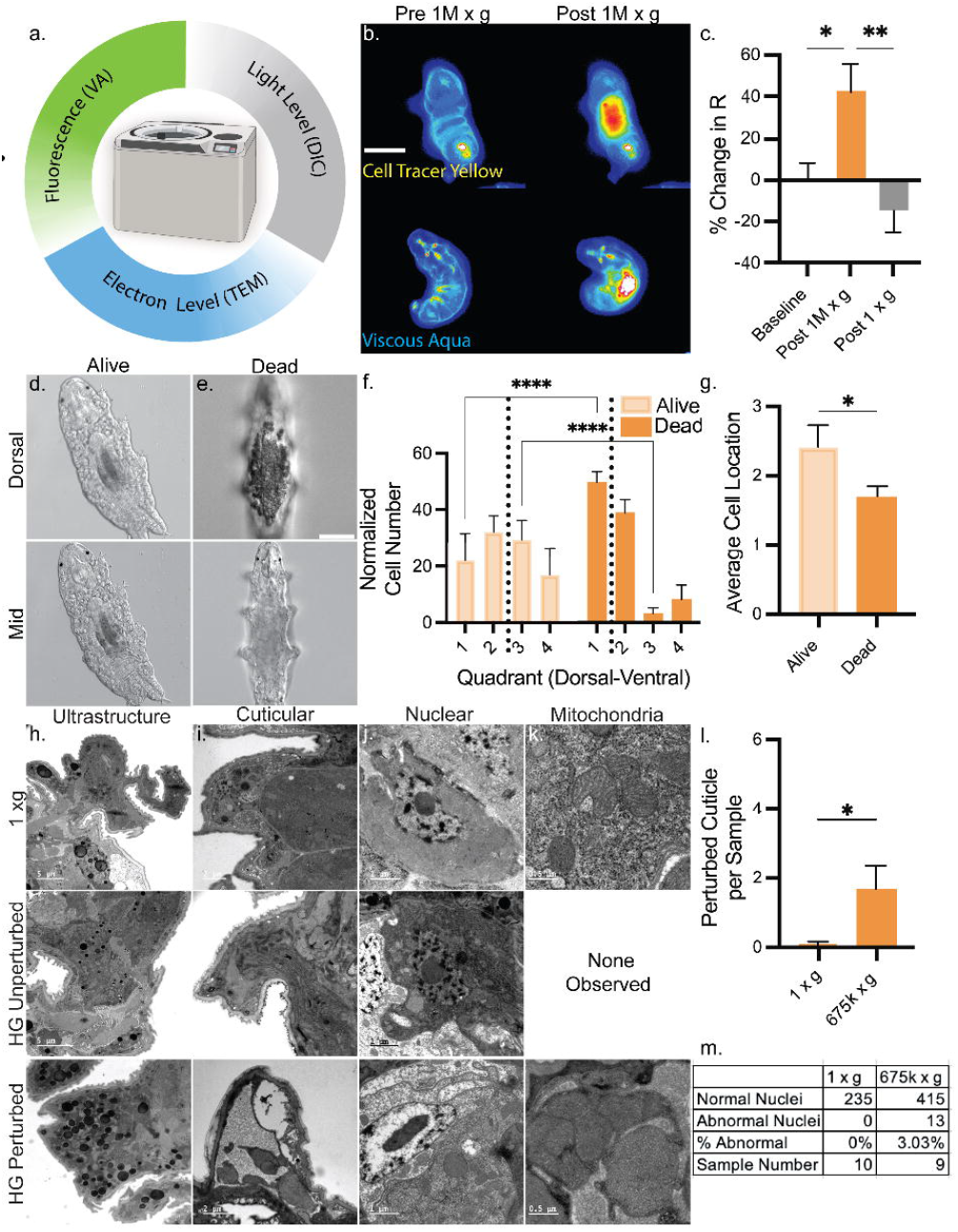
ROS production is required for entry into osmotic stress tun and survival in anoxia and HG. a) Schema depicting the functional activation of DCF-DA, by intracellular esterase uncapping and reactive oxygen species oxidation to increase fluorescence. b) Series of micrographs of tardigrades stained with DCF-DA taken 10 minutes after recovery from extreme stress states. Scale is 50 μm. c) Quantification of fold changes in fluorescence intensity of DCF-DA staining before and after stress exposure, **, p= 0.0079, ***, p= 0.0002, ****, p <0.0001. d) Survival rates of tardigrades exposed to extreme stressors in the presence of 20 mM cell-permeable glutathione, ****, p < 0.001 and *, p = 0.0486. e) Survival rates of animals pretreated with NEM, prior to exposure to extreme stressors, **, p=0.0071. f) Schematic of NOX 5-based ROS production and functional blockade of FAD cofactor catalysis by DPI. g) Survival rate of animals exposed to extreme stressors in the presence of NADPH oxidase Inhibitor, DPI, **, p = 0.0079 and *, p = 0.0191.

To investigate whether this spike in ROS might contribute to the mechanisms of resilience to HG, we inhibited intracellular ROS production with cell-permeable glutathione methyl ester (see Materials and Methods). We found this reductant led to a marked (57%) decrease (p=0.0486) in survival following 10 min centrifugation and a 16-hour recovery period (Fig. 5d), suggesting that ROS may be required to promote an HG-resistant state. To test whether ROS might do so by causing disulfide bond formation, we analyzed the effect of pretreating the animals with N-ethylmaleimide (NEM), which blocks free thiols in the cell. We assessed the efficacy of NEM in blocking free thiols by subsequently staining with equimolar Fluorescein-5-NEM adduct, which revealed an apparent ∼50% reduction in thiols in the NEM-treated animals compared to controls (Fig. S15). Despite partial reduction of free thiols, we found that NEM did not alter survival following exposure to HG (Fig. 5e), suggesting that the ROS-dependent process that protects animals from this extreme physical stress may not involve disulfide bond formation.

We found that ROS production, and its requirement in resilience to stress, extends to osmotic and anoxic stress: we observed a 3.6-fold increase in the intensity of DCF-DA staining as a result of osmotic stress (p=0.0079) and a 4.5-fold increase in response to CO_2_-sparged water (p=0.0002), similar to the elevation of ROS observed following exposure to HG (Fig. 5b and c). Further, we found that pharmacological reduction of ROS activity by glutathione inhibited entry into the tun state in response to osmotic stress (Fig. 5d and Fig. S16) and led to a 95% reduction in survival of animals following recovery from anoxia (Fig. 5d, p<0.0001). While pretreatment with NEM prior to the anoxic stress did not alter survival, we found that survival to osmotic stress was decreased by 38% (Fig. 5e, p=0.0071), suggesting that resilience to this osmotic stress (as previously seen (13)), but not anoxia or HG, requires disulfide bond formation.

We next sought to identify the cellular source of ROS that is produced in response to these three stresses. A primary site of ROS production is from the reaction catalyzed by NADPH oxidase (NOX), which produces superoxide at the extracellular surface. The superoxide, in turn, is converted to H_2_O_2_ prior to entry into the cell via diffusion (45). In contrast to the seven NADPH oxidase isoforms in mammals, the tardigrade genome encodes exclusively one, NOX-5, a simplified version of NADPH oxidase that uses calcium regulation and FAD-dependent catalysis to convert molecular oxygen to superoxide (45) (Fig. 5f). The NOX-5 gene has been duplicated five times in the tardigrade genome, suggesting its potential importance as a regulator of tardigrade physiology (10), (Fig. 5f). We found that inhibition of ROS production from NADPH oxidase by diphenyleneiodonium chloride (DPI; see Materials and Methods) significantly decreased survival following both osmotic stress (50% decrease, p=0.0079) and HG (69% decrease, p=0.0191) but had no significant impact on the survival of animals that experienced anoxia (CO_2_ sparging, Fig. 5g). These findings suggest that NADPH oxidase serves as a source of ROS that promotes survival for some, but not all, extreme stress conditions.

## Discussion

Life forms have evolved to survive in a broad range of environments on this planet. These adaptations can lead to states that allow organisms to withstand extraordinary conditions, including those that no organism has ever experienced over the 3.5+ billion years of evolution. We have found that such secondary effects of adaptation extend to the extreme physical stress of 1 million x *g* of simulated HG. While this extreme HG would be expected to stratify cellular components of different densities, we found no evidence of density-dependent stratification of cellular components, potentially owing to the measured increase in intracellular macro- and micro-viscosity observed across all extreme stressors explored in this study. Extending recent findings implicating ROS production as a major contributor to establishing cellular resilience in tardigrades, we found that ROS is required for survival or entry into all quiescent states tested. This ROS originates from NADPH oxidase under osmotic stress and HG exposure. We postulate that quiescent states, regulated by ROS arising from NADPH oxidase, reflect decreased molecular motion and dramatic increases in intracellular viscosity, rendering the animals resilient to extreme physical force experienced at 1 million times the force of Earth’s gravity.

As others have proposed (18), we found that intracellular viscosity significantly increases in response to hyperosmotic stress (Fig. 1h), likely owing to protein-mediated gelation induced by water loss. The enhanced sequestration suggested by the reduced mobile fraction of mitochondria indicates that the gelation matrix is compact, capable of restricting the overall motion of organelles suspended in the cytoplasm. Microviscosity was also observed to be increased under stress. At this scale, micro-diffusivity, influenced by mesh size, controls how molecules move; therefore, mesh size must be no greater than an order of magnitude larger than the dye molecule to affect microviscosity measurements (46). In the case of a small molecular rotor dye, such as Viscous Aqua (approximately 10-20 Å in diameter), our findings suggest a pore size of no greater than 10-20 nm within the cytosol (47); such a conclusion is consistent with the measured pore size in isolated fibers of CAHS D, a TDP, under desiccating conditions (18).

We had predicted that viscosity might decrease during anoxic conditions, as previous studies had shown cellular volume increases, potentially resulting from water uptake (8). However, in contrast, we found that intracellular viscosity increased significantly in anoxic animals, characteristic of a viscous fluid rather than a hydrogel structure, as organelle mobility was retained in these stressed states. Further, the kinetics of return to normal viscosity were rapid (<10 mins), supporting the hypothesis that a canonical hydrogel structure, which may take longer to resolve, does not form under these conditions. These findings suggest the existence of a distinct cytosolic viscosity-altering mechanism that is independent of water loss and that exhibits different kinetics of resolution.

To provide context for the magnitude of the viscosity change we measured (up to 125 cP under stressed conditions), the cytoplasm of certain human cancer cell lines ranges from 1.93 cP to 3.72 cP (48) and up to 15 cP in ametabolic *Talaromyces macrosporus* ascospores (49). The observed disparity between our datasets (8 cP cytoplasmic viscosity in active tardigrades vs. 1-3 cP cytoplasmic viscosity in other cells) may result from interactions of the mitochondria with other cellular components. The expression of Genetically Encoded Multimeric Particles (GEMs) or intracellular injection of microbeads was not feasible in the context of this study, owing to physical limitations of single cell microinjections and lack of stable transgenesis in these animals. As such it is likely that future studies may further refine our findings. It is important to note that the 15-minute time scale is likely too brief to allow for large-scale (23% change in mobile fraction; Fig. 1f) stress-induced hyperfusion of the mitochondrial network, which requires 2-3 hours in cultured cells (50), indicating that mitochondrial fusion dynamics are likely not responsible for the observed viscosity change.

The observed viscosity changes at the organismal and cellular level might underlie resilience to extreme physical force. Previous studies reported some lethality of tardigrades exposed to only <2% of the gravitational force (16,060 x g) we observed full survival (38). However, we note that our study was performed in an ultracentrifuge under vacuum and temperature control, which prevents fluctuations resulting from aerodynamic heating that could have killed the animals in the previous study. In comparison, we found that *C. elegans*, which was reported to survive 400,000 x g (37), is unable to survive 1M x *g* as a result of large-scale loss of cellular integrity of the animal (Fig. 3 c and d).

It is important to note that, while in nature tardigrades have never experienced the extreme HG used in this study and would not have been subjected to selective pressure to adapt to this extreme physical force, their resilience to it likely reflects cross-tolerance (51) arising from adaptations to other stressors, such as osmotic stress. Intriguingly, the survival rate of *H. exemplaris* under HG increased in high viscosity stress states, especially hyper-osmotic stress (enhanced survival by 2.66-fold longer), suggesting that cytosolic viscosity changes and molecular sequestration enhance survival to HG. Consistent with this hypothesis, tardigrades exposed to HG maintain elevated micro-viscosity fully ten minutes after returning to normal gravity and return to normal viscosity with similar kinetics to those seen after removal from hyperosmotic stress.

Consistent with the current model of desiccation and osmotic stress, we find that ROS production is necessary for survival across all tested states. We expanded upon this model with evidence suggesting that intracellular ROS production via NADPH oxidase is required for survival in response to both HG and hyperosmotic stress. We note that DPI may have other flavoenzyme targets; thus, the requirement for NOX-5 will need to be evaluate further in future studies using, for example, RNAi knockdown (52). Regardless, our findings support the view that NADPH oxidase may be a key regulator of osmotic stress response, acting as a primary source of ROS.

Our findings of *in vivo* cytosolic viscosity changes across multiple stress states in tardigrades suggest two distinct changes in response to different stressors: a viscous and sequestering “hydrogel-like” structure during osmotic stress, and a viscous non-sequestering fluid-like cytosol in anoxic animals. We suggest that HG induces a high-viscosity state that is akin to that of the hyperosmotic stress response, inducing viscosity changes of similar magnitude, kinetics of resolution, and reliance upon NADPH-oxidase mediated ROS signaling. These high viscosity states are likely to account for the startling resilience of these animals to extraordinary HG force of 1M x g, which, to our knowledge, makes them the most HG-resilient animal to date.

## Acknowledgments

We appreciate the NRI Microscopy Facility (UC Santa Barbara) for providing microscopes, UC Berkeley Electron Microscopy Lab for TEM imaging support, Zvonimir Dogic’s laboratory for access to their ultracentrifuge, and ThermoFisher Scientific’s applications team for help with freezing methods. This work was supported by NIH grant 5R01HD081266 and the Wilcox Family Chair in Biotechnology to J.H.R. M.J.K. was supported by NIH award 1 F32 AG081056.

