## Supplemental Information for "Evidence that local viscosity and NOX-dependent ROS increases render the tardigrade H. exemplaris resilient to extreme physical force"

**This PDF file includes:**

Methods Section
Figures S1 to S16
Tables S1 to S3
SI References

**Supporting Information Text**

**Materials and Methods:**

*Tardigrade, Algae and C. elegans Culture Maintenance:*

Tardigrades were cultured under the conditions described by (1, 2). Briefly, cultures of *Chlorococcum hypnosporum* (Carolina, 152091) were grown in 500 mL glass Erlenmeyer flasks containing 30 mL of Bold’s Basal Medium, 25%V/V autoclaved soil extract, and 120 mL sterile spring water to a total volume of 150 mL (Nestle Pure Life, 44221229). Cultures were grown at 20 °C, with shaking and constant light, for 3-5 weeks prior to use. Tardigrade (Sciento, Z151) cultures were maintained in sterile 35 mm plates containing 3.2 mL of sterile spring water and 0.8 mL of algal culture. Cultures were fed fresh algae weekly, washed biweekly. Cultures were maintained at room temperature (22-23 °C) with a 10-hour to 14-hour light-dark cycle. C. elegans worms were grown and maintained on OP50 E.coli at room temperature (23◦C) using standard methods(3). Dauer formation was induced by overcrowding, which occurred 8–12 days after chunking onto seeded NGM plates. Dauer larvae were isolated from 1 to 5 starved NGM plates using 1% SDS and a 30% sucrose gradient as described previously(4).

*Anaerobic environment creation and validation:*

990 µL of spring water containing (10mM potassium phosphate (pH 7.5 +/- 0.03) 20 mM DL Lactate was sparged in a Serum Vial (PATIKIL, Amazon) sealed with 13 mm Butyl stoppers (PATIKIL, Amazon) secured with 13mm aluminum caps (PATIKIL, Amazon), using high purity nitrogen (Airgas, UN1066) for 15 minutes before addition of ~~the addition~~ 10 µL ~~of~~ Oxyrase (Oxyrase EC-0005)(5). The vials ~~was~~ were incubated at 37°C for 40 minutes. Animals were then placed in a 20 µL droplet of water at the bottom of a clean serum vial and transferred into the glove box. Vials were then capped in the glove box, and 200µL of Oxyrase-treated anoxic spring water was added to each sealed vial. D.O. measurements were made across two independent experimental days across six independent vials, using a needle-type oxygen microsensor (Presens, NTH-PST7) and Fibox 4 Trace (Presens) D.O. meter

*FRAP Imaging:*

30 mixed-stage tardigrades were isolated from culture prior to the addition of 20 µM Mito tracker Deep Red Dye (1 mM water stock, stored at -20 °C). Dye incubation was performed at room temperature in the dark for one hour prior to washing with 1 mL of spring water. Animals were then imaged on a Leica SP8 confocal microscope using a HC PL APO CS2 63x/1.30 glycerol immersion objective. Tardigrades were placed in spring water (pH 7.0), 0.5M Mannitol (pH, 7.0), $CO_2$ sparged spring water (0.045 +/- 0.05 ppm D.O., pH 3.4), $N_2$ sparged Oxyrase sodium lactate buffer (0.05 +/- 0.028 ppm D.O., pH 7.0) or normoxic sodium lactate buffer (8ppm D.O., pH 7.0) for 10 minutes prior to FRAP imaging. Animals were placed on a 3% agar pad secured with a no. 1.5 glass coverslip, in the presence of polystyrene beads (Polyscience, 00876-15, 0.1µm in diameter) to limit animal movement. A 2.5 µm in diameter circular region of the animal’s head was bleached for 0.65 seconds using a combined 675 and 594 laser line at full power, and recovery of Mito Tracker Deep Red fluorescence was monitored for the subsequent 500 frames. Data were analyzed using easy-FRAP(6) for circular bleach patterns or SIM-FRAP(7) for square bleach patterns, exponentially fit with an $R^2$ of 0.7 or higher. Mobile fraction was calculated via the following equation.

$$M_f = \frac{(F_f - F_0)}{(F_i - F_0)} \qquad (1)$$

Where $F_f$ is final fluorescence, $F_o$ is fluorescence postbleach, and $F_i$ is the initial fluorescence of the bleached region. $t_{1/2}$ was then used to calculate the diffusion coefficient using a Soumpasis model(8). The Soumpasis model is as follows:

$$D = \frac{0.224 r_n^2}{\tau_{\frac{1}{2}}} \qquad (2)$$

Where $D$ is the apparent diffusion coefficient, $r_n^2$ is the radius of the bleached region, and t½ is the time at half maximum recovery. Einstein-Stokes Equation, was used to estimate the viscosity experienced by a mitochondrion of 0.5 µm x 0.5 µm x 1 µm, modeled as a three-dimensional tumbling ellipse (9).

$$D = \frac{k_B \cdot T}{\xi} \; where \; \xi = \frac{6\pi \cdot \eta \cdot a}{ln\, ln\, \left(2 \cdot \frac{a}{b}\right)} \qquad (3)$$

Where $D$ is the diffusion coefficient, $k_B$ Is the Boltzmann constant, $T$ is temperature in Kelvin, $\xi$ is the drag coefficient, and $\eta$ is viscosity, and $a$ is the major axis of the ellipsoid, and $b$ is the minor axis of the ellipsoid.

*Viscous Aqua and DCFDA Imaging:*

Dye stocks of Viscous Aqua (2.5 mg/mL, VA; Ursa Bio Sci. 140082) and CTY (5 mM, Thermofisher, C34573) in DMSO were maintained at -80 °C. 30 mixed-stage tardigrades were incubated for 1 hour at 21-23 °C in the dark in a solution either 96 ng/ul of VA, 5 µM CTY, and 0.65% DMSO or 20µM DCFDA prior to washing with 1 mL of spring water. All animals were imaged within a 2-hours of staining. Imaging was performed on a Nikon TI-Eclipse equipped with an Orca Flash 2.8 (Hamamatsu) sensor, using TRITC (Semrock, TRICTC-B-NTE-ZERO) and CFP (Semrock, CFP-2432C-NTE-ZERO). Animals were placed on a glass slide, and images were taken before, 10 minutes after exposure, and 10 minutes after return to ambient environmental conditions. Whole animal fluorescence intensity measurements of both VA and CTY were obtained via FIJI and presented as %ΔR/R values calculated using,

$$\% \frac{\Delta R}{R} = \frac{R - R_o}{R_o} \; where \; R = \frac{Viscous\ Aqua\ Fluor.Int.}{Cell\ Tracer\ Yellow\ Fluor.Int.} \qquad (4)$$

*Spectroscopy and anoxic spectroscopy:*

Samples of VA dye (25 µg/mL) were diluted in normoxic spring water (8ppm, pH 7.0), nitrogen and Oxyrase-treated spring water (0.17 +/- 0.01 ppm, pH 7.0), and CO2-sparged spring water, (0.045 +/- 0.05 ppm D.O. ppm D.O.) (Fisher, 12565501) and covered with 100 µL of either normoxic mineral oil (Sigma, H8898) or nitrogen-sparged mineral oil. Plates were then sealed with a plate seal (Thermo Scientific, AB-0558) under normoxic or anoxic conditions (0.13 +/- 0.1 ppm oxygen in a nitrogen-charged glove bag). Samples were then measured on a TECAN Infinite 200 pro plate reader, and fluorescence intensity was measured following 25 flashes at 430 nm +/- 9nm excitation, with emission collected from 490 nm +/- 20 nm. Data are reported as background (empty well) subtracted fluorescence intensity counts for all conditions. Data were collected in quadruplet well measurements, in triplicate, for each of three independent experimental days.

*Centrifugation:*

Six mixed-stage tardigrades were isolated from culture and placed in an ultracentrifuge tube (Thermofisher ,45237) containing 1mL of spring water. Centrifuge tubes were then balanced to a weight of 2.65 +/- 0.02 grams and centrifuged (Sorvall MTX 150 Micro Ultracentrifuge, Thermofisher), in a S140-AT rotor (Thermofisher) at an max RCF of either 100k, 675k or 1M x g for 10 minutes, or 1M x g for 1, 10, 20, 30 and 45 minutes not including ramp, which was approximately 2 minutes in duration. Following centrifugation, each sample was dispensed into a fresh 96 well-plate assay plate (Fisher, 12565501), alongside an unspun control cohort of the same size. All tardigrades were then monitored for time to first movement, time to coordinated movement, survival, egg-laying, and progeny production for the subsequent 15-16 days.

*Progeny and Survival Rate Assay:*

Tardigrades from the centrifugation experiment were subsequently monitored every 2-3 days for survival rate (determined by coordinated locomotion), egg production, and reproductive success (i.e., number of viable progeny). During each monitoring session, tardigrades from each condition were propagated to the next well in a 96-well plate to prevent the mixing of parents and progeny. These three measurements were obtained across at least three individual experiments and were monitored every Monday, Wednesday, and Friday for 15-16 days following centrifugation.

*Light level examination of storage cell localization post centrifugation:*

Animals exposed to HG for 30 min at 675,000x g were monitored after exposure and sorted into mobile and non-mobile cohorts. Images were obtained (20 minutes following exposure) under 20x DIC microscopy on a Nikon Ti microscope, performing a z-stack from the ventral to dorsal aspect. The z-stack was divided into four equal parts, and the first section from each quartile was used for manual counting of storage cell numbers. Each condition contained at least four individual animals.

*Freezing and TEM:*

Ultracentrifuge tubes were filled with 1mL of Spring Water, and a 40 mm piece of twine (Home Depot, #70077). The tube weight was adjusted to 2.65 +/- 0.02 g and frozen at -20°C. The ice was then removed from the tube using the twine, and a sample containing 100 tardigrades in 100 µL of water was added to the bottom of the tube. The ice was quickly replaced and allowed to settle, leaving a small pocket of liquid at the bottom of the ice. Samples were then immediately placed into a prechilled rotor (S140-AT, -20°C) vacuum chamber at (0°C), and the vacuum was applied. The sample was held at 675,000 x g for 3 minutes. Samples were immediately removed and fixed in 3% Glutaraldehyde and 4% PFA, and stored at 4 °C. imaged at 20x DIC using a Nikon Ti-Eclipse equipped with an Orca Flash 2.8 Camera (Hamamatsu). Measurements were taken for dorsal-ventral, head-tail, and the point of longest width using a FIJI macro. From these measurements, the volume of an oblong shape was calculated as an estimate of total animal volume. Samples frozen and fixed under HG were then stored in fixative for 1 week prior to tissue preparation. Samples were embedded into 2% agarose (EMS, Agarose SFR™ 10207) and fixed overnight in 3% PFA and 4% Glutaraldehyde. Samples were washed 3x for 15’ in the PhEM buffer and postfixed in 0.5% Osmium tetroxide in PhEM (EMS) and dehydrated in an acetone series (30%, 50%, 70%, 85%, 95%, and 3x 100%) for 15 minutes each. Tardigrades were then embedded in Epon Resin (EMS, Cat #: 14120) over 5 days and baked for 2 days at 60°C, prior to ultrathin sectioning on a Leica EM UM6 Ultramicrotome. Post-staining with Lead Citrate and Uranyl Acetate was followed by imaging on a Technai 12 TEM.

*Drug Treatments:*

Drugs were stored at ~~at~~ -20 °C in 30 mM - 5 mM stocks in water or DMSO, except NEM, which was made fresh. Final stock concentrations were diluted from stock in spring water at pH 7.0: Glutathione-monoethyl ester 20mM (GSH, Milpore Sigma), N-ethylmaleimide 30 µM (NEM, Pierce), and diphenyleneiodonium chloride 5µM (DPI, Cayman Chemical). A cohort of 6 animals ~~were~~ was pretreated for 1 hour prior to exposure to extreme environments (O/N spring water control, 2 hours mannitol, O/N Anoxia, and 10 minutes HG). Following exposure, animals were washed 3x with 400 µl of water and allowed to recover in the absence of food for 24 hours before survival was assessed based on locomotion.

*Propidium iodide Dye Loading Assay:*

To assess the function of propidium iodide as a measure of cell survival post-centrifugation, we isolated 20 tardigrades from cultures and washed them three times in 1 mL of spring water. We then placed them into 1.5 mL Eppendorf tubes containing 100 µL of spring water.~~, to the tube along with the tardigrades.~~ Two of the Eppendorf tubes were then closed and incubated at 45 °C for one hour to ensure tardigrade death; the remaining tubes were maintained at room temperature with their lids open. Following heat shock, Propidium Iodide (1mg/mL, Sigma P-4170) and Hoechst were added at 1:100 to one heat-shocked and one live sample, and the samples were incubated for 30 minutes at room temperature in the dark. The remaining two samples were unlabeled with dye and served as controls for autofluorescence. Following dye loading, both live samples were treated with 5 µl of 10 mM Levamisole in water, and all samples were mounted onto agar pads and covered with a glass coverslip, with care taken to place spacers of clay to prevent direct contact between the glass and the agar. Imaging was performed on a Nikon Ti inverted epifluorescence microscope equipped with DIC optics. Images were obtained with an Orca-Flash camera set to 10 ms exposures for DIC, and 500 ms exposures for the Texas Red channel (Semrock, TRITC-B-NTE-ZERO) and the DAPI Channel (Semrock, DAPI-1160B-NTE-ZERO). Images were focused in each channel to obtain the clearest image. Images were then analyzed in FIJI/ImageJ, quantifying whole-animal propidium iodide fluorescence intensity. Values were normalized to the average background fluorescence of live and dead samples.

*Modeling of density-dependent stratification:*

We developed a customized numerical simulation tool using the finite element software COMSOL Multiphysics (www.comsol.com) to simulate the deformation of a cell and the underlying protein distribution within the cytoplasm under extreme *g*-forces approaching 500,000 times Earth's gravity. Here we model the cell's cytoplasm as an 8 $\mu$m diameter sphere containing a Newtonian fluid. The surrounding cell wall membrane is modeled as a thin 7 nm thick linear elastic solid. For simplicity, the simulation is conducted using a 2-D axisymmetric coordinate geometry. The fluid motion in the cytoplasm is simulated using the incompressible Navier-Stokes equations, while the cell wall membrane is simulated using Hook's law and the Cauchy stress tensor. An extreme *g*-force of 500,000 times gravity is applied to the cell, which causes the cell to deform over time, as it is pushed against a flat rigid surface. The deformation of the cell is simulated by mapping the cell geometry into a deformed coordinate system, using a Yeah mesh smoother. As the cell body undergoes deformation due to extreme *g*-force, we simultaneously simulate the volume fraction of protein concentration within the cytoplasm. Protein distribution is modeled using the conservative scalar equation

$$\frac{\partial \varphi}{\partial t} + (\mathbf{u}\cdot\nabla)\varphi = \nabla\cdot\left(D\nabla\varphi + v_p\varphi\right) \qquad (5)$$

where $\boldsymbol{\varphi}$ describes the volume fraction of protein, **u** is the fluid velocity vector of the cytoplasm, D is the protein diffusivity, and $v_p$ is the velocity of the protein relative to the cytoplasm due to buoyancy under extreme G-force. The cytoplasm viscosity is a function of protein concentration. Following Krieger and Dougherty (1950), we model the dynamic viscosity of the cytoplasm as

$$\mu = \mu_0\left(1 - \frac{\varphi}{\varphi_m}\right)^{-n\varphi_m} \qquad (7)$$

where $\mu$ is the dynamic viscosity, $m_0$ is the viscosity without proteins, and $j_m$ is the maximum volume fraction due to steric interactions. Krieger and Dougherty (1950) report the exponent n = 2.5. We model the diffusivity to decrease with increasing viscosity using

$$D = D_0\left(1 - \frac{\varphi}{\varphi_m}\right)^{n\varphi_m} \qquad (8)$$

where $D_0$ is the diluted protein diffusivity. In the numerical simulation effective gravity is ramped up at a rate of 500,000 *g*-force for a time period of 200 seconds, which corresponds to the typical ramp up time for our centrifuge. The initial protein volume fraction is set to $\boldsymbol{\phi}_0$ = 0.2. The deformed cell and protein concentration are shown in the time in Figure S11. The geometry is axisymmetric, where the horizontal axis is the radius, *r,* and the body is revolved around the vertical *z*-axis.

*NEM Blockade and Survival:*

To assess that NEM blocked available thiol groups in the Tardigrade, animals were pretreated with 30 μM NEM (for one hour at room temperature prior to treatment with 30 μM NEM-dye adduct Fluorescein-5-Maleimide (Cayman Chemicals, CAS 75350-46-8) for 30 minutes. Animals were then washed 3 times with 100 μL of water prior to mounting on 3% agar pads and epifluorescence microscopy using a Nikon TI-Eclipse equipped with an Orca Flash 2.8 (Hamamatsu) sensor, and FITC Filter(FITC-B-NTE-ZERO).

Relative fluorescence intensity was monitored using FIJI/ImageJ -and reported as a fold change relative to the non-drug-treated control.

**Figures:**

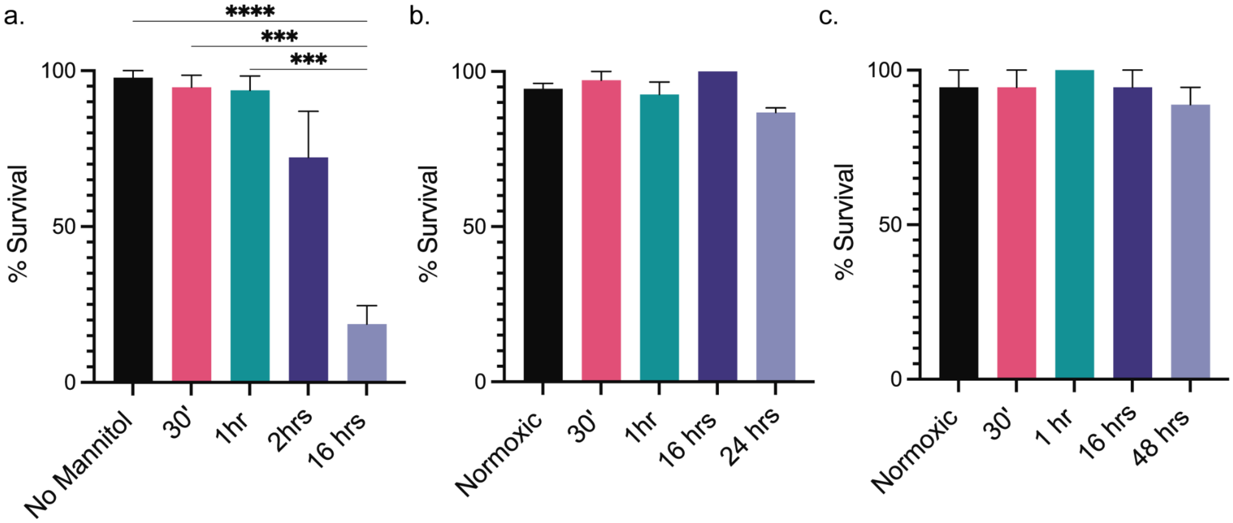


**Figure S1: Survival rates under osmotic shock, CO2 and $N_2$-sparged anoxic conditions, .** a. Survival rates of animals exposed to osmotic shock 0.5 M mannitol for 30', 1hour, 2hours, and 16 hrs. Kruskal-Wallis test with Dunn's multiple comparisons test, ****, p>0.0001, and ***, p = 0.0003 and 0.0004. b. Survival rate for animals exposed to $CO_2$ sparged spring water 0.045 +/- 0.05 ppm D.O. 30' 1 hour, 16 hours, 24 hours. Kruskal-Wallis test with Dunn's multiple comparisons test revealed no significant differences. c. Survival rate for animals exposed to $N_2$ sparged and Oxyrase treated spring water 0.05 +/- 0.028 ppm D.O. for 30', 1 hour, 16 hours and 48 hours. Kruskal-Wallis test with Dunn's multiple comparisons test revealed no significant differences. Data in all panels represent the mean ± S.E.M. survival rate across 6 animals per trial and 9 trials.

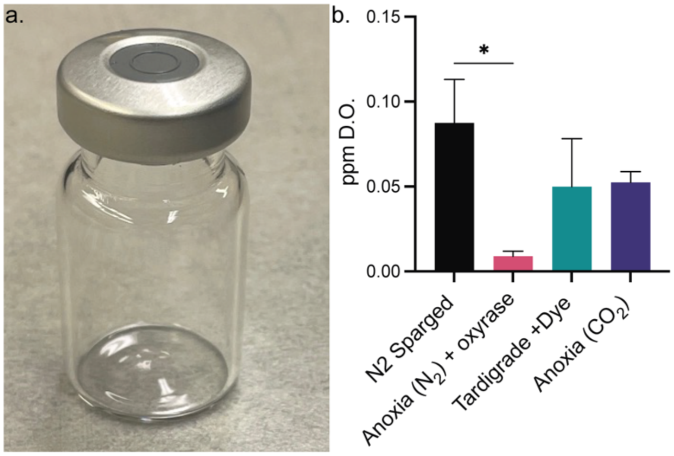


**Figure S2: Measurement of D.O. levels attained under Oxyrase-treated vials and survival rates under anoxic conditions.** a. Image of 2mL serum vials sealed with a rubber self-sealing cap and metal clamp ring. b. D.O. measurements obtained from anoxic vials sparged with ultrapure nitrogen and treated with Oxyrase (black), nitrogen sparging only(pink), and prestained tardigrades added to nitrogen-sparged and Oxyrase-treated vials. Data represent the mean and S.E.M. of individual measurements obtained from 4-9 individual vials. Kruskal-Wallis test with Dunn's multiple comparisons test revealed no significant differences

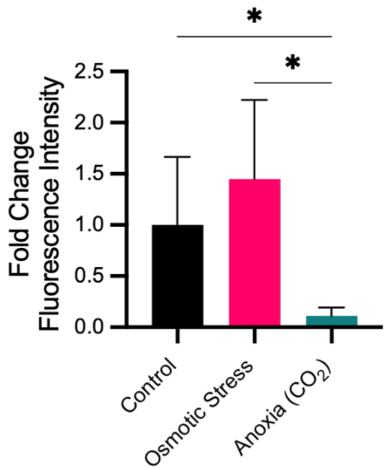


**Figure S3: Mitotracker ~~deep~~ Deep ~~red~~ Red fluorescence intensity under various environmental extremes.** Initial fluorescence intensity measurements of FRAP bleach ROI following exposure to control spring water, 0.5M mannitol, and 0.045 +/- 0.05 ppm D.O. $CO_2$ sparged water for 10 minutes.

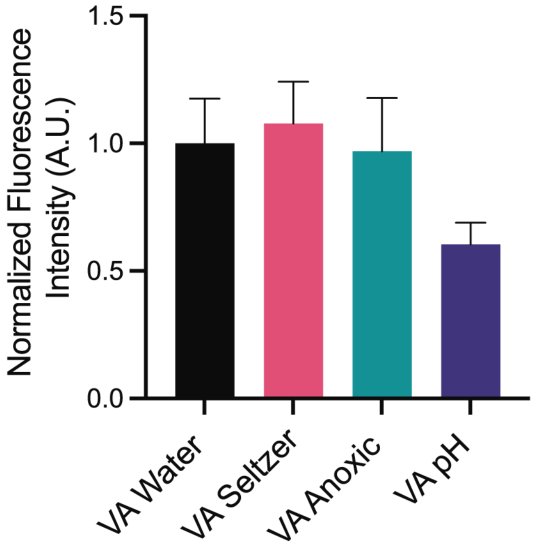


**Figure S4: Anoxic and Hypoxic Spectroscopy of Viscous Aqua.** Spectroscopic plate reader measurements of 25 μg/mL) VA under normoxic, anoxic, hypoxic, and low pH (pH 3, Citrate Buffer) conditions, reported as normalized fluorescence intensity from 6-9 individual wells across at least two independent experimental days. One-Way ANOVA, with Tukey's Multiple Comparisons Test, revealed no significant difference.

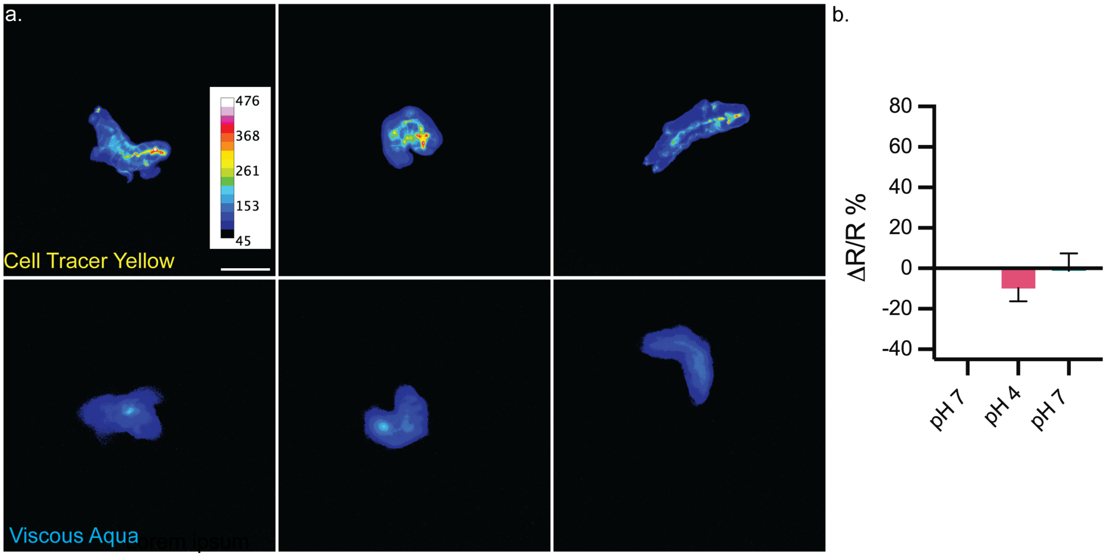


**Figure S5: Low pH does not alter cellular viscosity as reported by Viscous Aqua.** a. Image panel of CTY (top), VA (bottom) fluorescence intensity measured before (left), during (middle), and after (right) exposure to low pH 47 mM citrate buffer at pH 3 for 10 minutes. Data is pseudo-colored using a 16-color lookup table, and the scale is 50 μm. b. % *Δ*R/R representing the change in the ratio of VA to CTY across an N of 9 individual animals obtained across three individual experimental days. Brown, Forsythe, and Welch ANOVA with Dunnett's T3 test found no statistical significance.

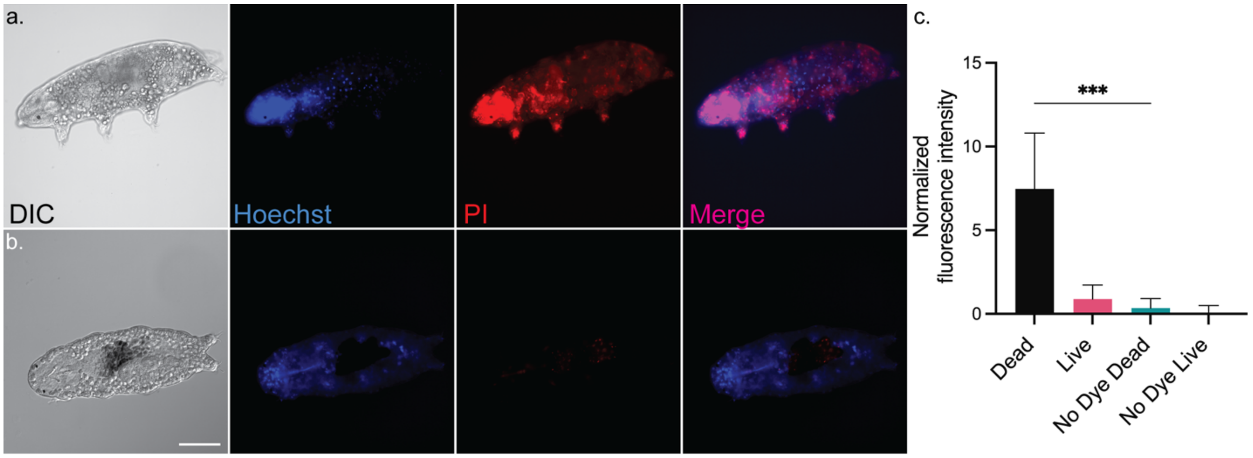


**Figure S6: Propidium iodide live-dead cell stain assay.** a) DIC, Hoechst nuclear counter stain, propidium Iodide, and fluorescence merged micrographs of dead Tardigrade exposed to heat and Propidium Iodide. b) DIC, Hoechst, propidium iodide, and fluorescence merged micrographs of active state animals exposed to PI, Hoechst, and Propidium Iodide at RT. The scale is 50 µm. c). Quantification of whole body normalized PI intensity in heat-treated (dead), non-heat-treated (live), and no PI dye-treated control animals. Brown Forsythe and Welch's ANOVA using Dunnett's T3 multiple comparisons, ***, $p<0.0006$.

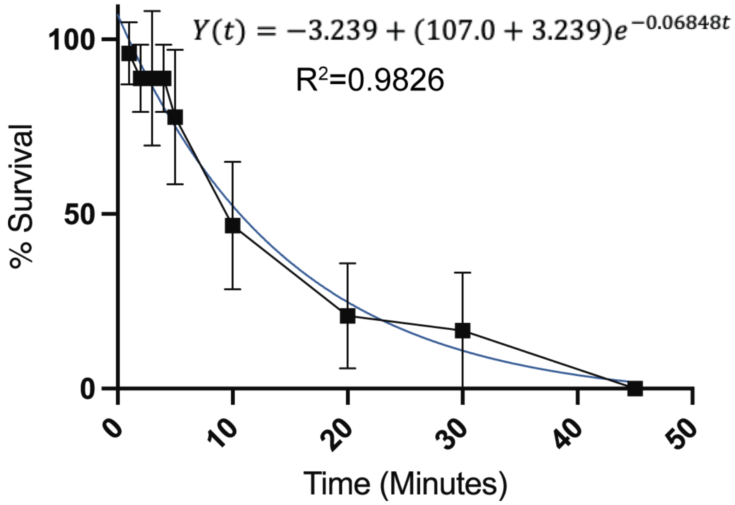


**Figure S7: Survival is exponentially related to exposure duration.** Percent survival following exposure to 1M x g for 0,1, 2, 3, 4, 5, 10, 20, 30 and 45 minutes. Data represents the average survival across at least 3 independent trials per time point, where each trial consisted of a cohort of 6 animals. Mean data was fit to a ~~singal~~single exponential decay function, whose equation is displayed above. Goodness of fit was determined by calculating the coefficient of determination ($R^2$) which was found to be 0.9826.

a.

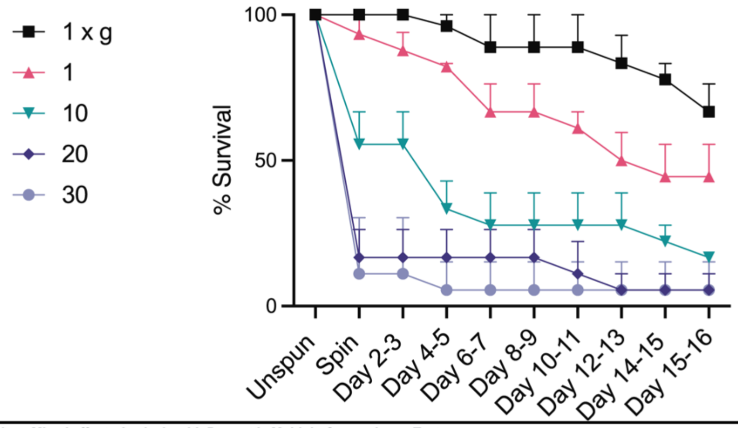


| Adjusted p-Values Mixed effects Analysis with Dunnett's Multiple Comparisons Test | | | | | | | | | | |
|---|---|---|---|---|---|---|---|---|---|---|
| | Unspun | Spin | Day 2-3 | Day 4-5 | Day 6-7 | Day 8-9 | Day 10-11 | Day 12-13 | Day 14-15 | Day 15-16 |
| 1 x g vs. 1 min | >0.9999 | 0.8477 | 0.4479 | 0.5622 | 0.1221 | 0.1221 | 0.0444 | 0.0049 | 0.0049 | 0.1225 |
| 1 x g vs. 10 min | >0.9999 | <0.0001 | <0.0001 | <0.0001 | <0.0001 | <0.0001 | <0.0001 | <0.0001 | <0.0001 | <0.0001 |
| 1 x g vs. 20 min | >0.9999 | <0.0001 | <0.0001 | <0.0001 | <0.0001 | <0.0001 | <0.0001 | <0.0001 | <0.0001 | <0.0001 |
| 1 x g vs. 30 min | >0.9999 | <0.0001 | <0.0001 | <0.0001 | <0.0001 | <0.0001 | <0.0001 | <0.0001 | <0.0001 | <0.0001 |
| 1 x g vs. 45 min | >0.9999 | <0.0001 | <0.0001 | <0.0001 | <0.0001 | <0.0001 | <0.0001 | <0.0001 | <0.0001 | <0.0001 |

b.

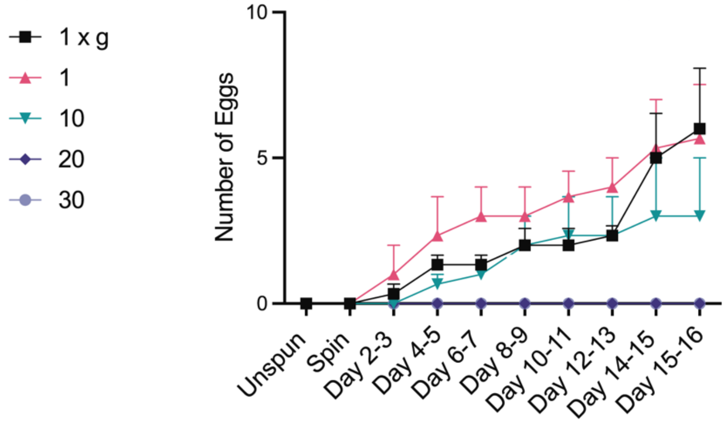


| Adjusted p-Values Mixed effects Analysis with Dunnett's Multiple Comparisons Test | | | | | | | | | | |
|---|---|---|---|---|---|---|---|---|---|---|
| | Unspun | Spin | Day 2-3 | Day 4-5 | Day 6-7 | Day 8-9 | Day 10-11 | Day 12-13 | Day 14-15 | Day 15-16 |
| 1 x g vs. 1 min | >0.9999 | >0.9999 | 0.9814 | 0.8653 | 0.4163 | 0.8033 | 0.3509 | 0.3568 | 0.9942 | 0.9993 |
| 1 x g vs. 10 min | >0.9999 | >0.9999 | 0.9927 | 0.9357 | 0.9969 | >0.9999 | 0.9951 | >0.9999 | 0.2684 | 0.0303 |
| 1 x g vs. 20 min | >0.9999 | >0.9999 | 0.9927 | 0.5532 | 0.583 | 0.2423 | 0.2617 | 0.1378 | <0.0001 | <0.0001 |
| 1 x g vs. 30 min | >0.9999 | >0.9999 | 0.9927 | 0.5532 | 0.583 | 0.2423 | 0.2617 | 0.1378 | <0.0001 | <0.0001 |
| 1 x g vs. 45 min | >0.9999 | >0.9999 | 0.9913 | 0.6764 | 0.7508 | 0.5295 | 0.6059 | 0.4736 | 0.0103 | 0.0011 |

c.

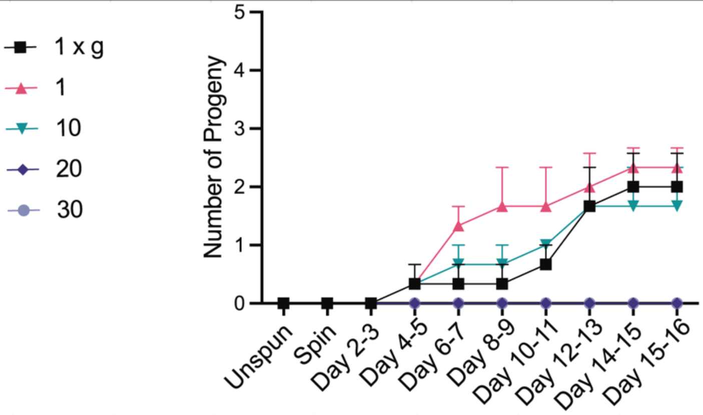


| Adjusted p-Values Mixed effects Analysis with Dunnett's Multiple Comparisons Test | | | | | | | | | | |
|---|---|---|---|---|---|---|---|---|---|---|
| | Unspun | Spin | Day 2-3 | Day 4-5 | Day 6-7 | Day 8-9 | Day 10-11 | Day 12-13 | Day 14-15 | Day 15-16 |
| 1 x g vs. 1 min | >0.9999 | >0.9999 | >0.9999 | >0.9999 | 0.1043 | 0.0147 | 0.1043 | 0.9191 | 0.9191 | 0.9191 |
| 1 x g vs. 10 min | >0.9999 | >0.9999 | >0.9999 | >0.9999 | 0.9191 | 0.9191 | 0.9191 | >0.9999 | 0.9191 | 0.9191 |
| 1 x g vs. 20 min | >0.9999 | >0.9999 | >0.9999 | 0.9191 | 0.9191 | 0.9191 | 0.4365 | 0.0013 | <0.0001 | <0.0001 |
| 1 x g vs. 30 min | >0.9999 | >0.9999 | >0.9999 | 0.9191 | 0.9191 | 0.9191 | 0.4365 | 0.0013 | <0.0001 | <0.0001 |
| 1 x g vs. 45 min | >0.9999 | >0.9999 | >0.9999 | 0.98 | 0.98 | 0.98 | 0.746 | 0.0388 | 0.0085 | 0.0085 |

**Figure S8: Survival, egg laying, and reproduction post centrifugation at 1M x for 1, 10, 20, and 30 minutes compared to a 1 g control group.** a) % Survival rate following 1M x g exposure, data represent the average and S.E. survival rate from 6 individuals, across three experimental days. b) Cumulative egg production following HG exposures, c) Cumulative progeny production following HG exposures. All error bars represent S.E. All statistical tests were performed with a mixed effects analysis and Dunnett's multiple comparisons test comparing each time point to 1 x g. Adjusted p-values are presented in the table below each respective graph.

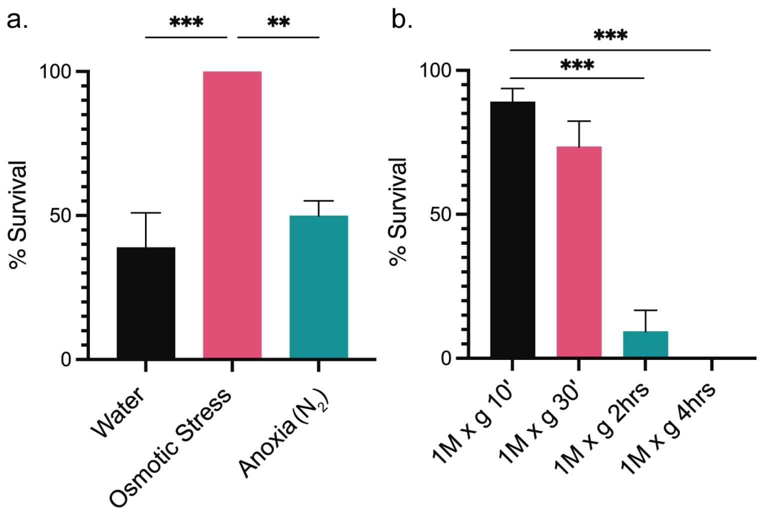


**Figure S9: Spinning under anoxic or osmotic stress.** a) Percent survival rates of animals exposed to 1M x g for 10 minutes under rearing conditions (black), 0.5 M mannitol (pink), and 0.5 +/- 0.1 atmospheric oxygen levels (green). Kruskal-Wallis test with Dunn's multiple comparisons test, ***, p =0.0004 and **, p = 0.0024. b) Percent survival rate of animals exposed to 1M x g in the presence of 0.5 M mannitol, for 10 minutes, 30 minutes, 2 hours, and 4 hours duration at max speed. Kruskal-Wallis test with Dunn's multiple comparisons test, ***, p = 0.0002 and 0.0007, respectively. All tests were performed across 7-12 trials, each containing 6x animals, and completed over three experimental days.

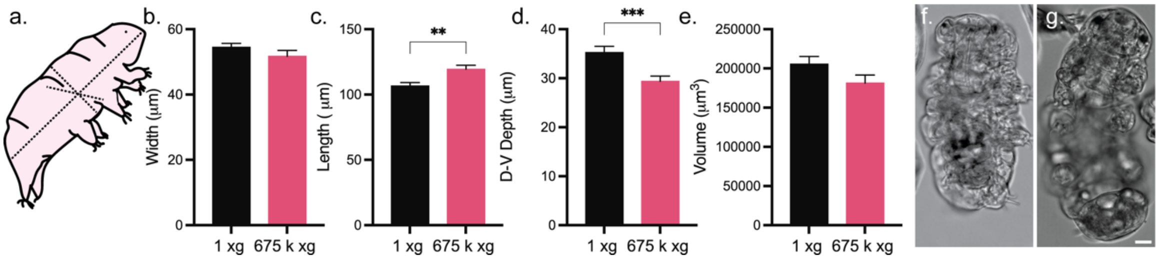


**Figure S10: Light level anatomy of tardigrades post HG exposure.** a) Schema of measurements, length, width, and depth obtained for quantification. b) Quantification of length changes in those exposed, and frozen and fixed under 675,000 x g, as compared to those frozen and fixed at 1x g. T-Test **, p =0.0021. c) Quantification width at the center of mass obtained from those exposed frozen and fixed under g-force. T-Test: no significant difference. d) Depth in µm measured as the z-stack step size times number of stack images from those frozen and fixed under g force, T-Test ***, p= 0.0007. DIC micrograph of Tardigrade frozen and fixed at 1x g (f), or frozen and fixed at 675,000 x g (g). The scale is 10 µm.

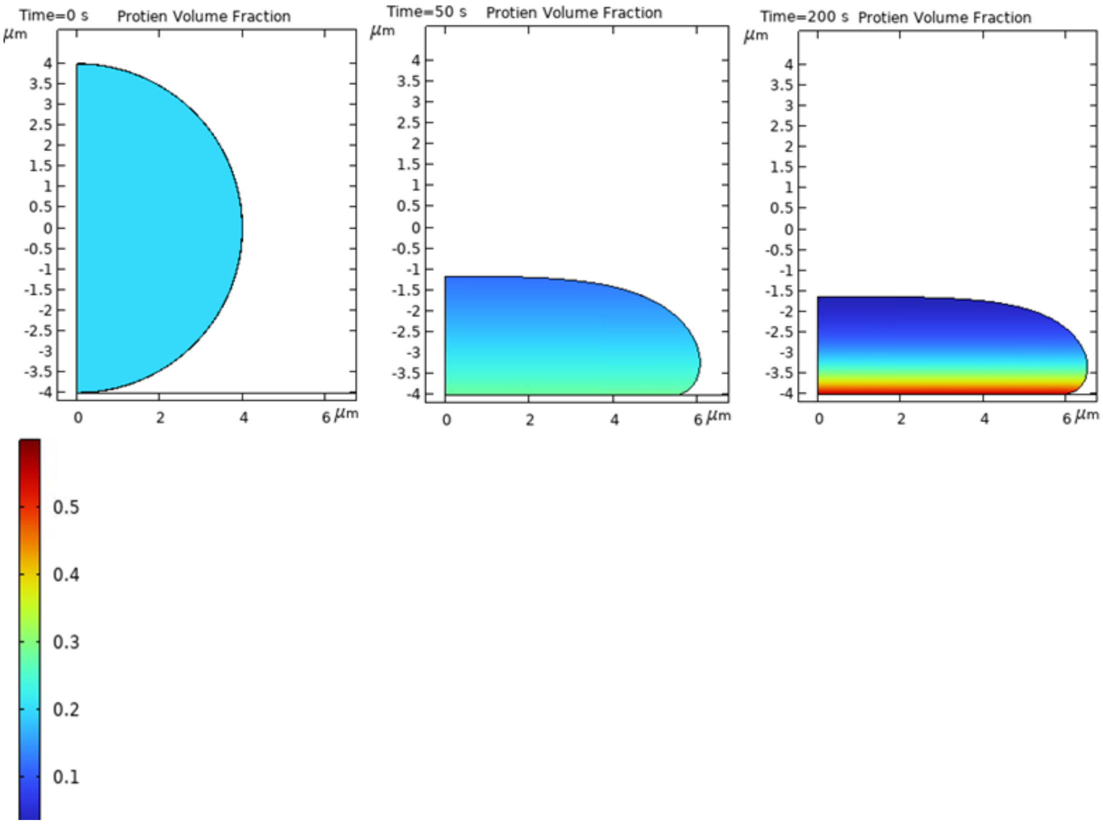


**Figure S11: Time sequences showing cell deformation and protein concentration for times *t* = 0, 50, and 200 seconds.** The protein volume fraction, $\phi$, is denoted by the color map shown to the right, which varies from $\phi$= 0 volume fraction (indicated by blue) to a maximum of $\phi_m$=0.6 (indicated by red).

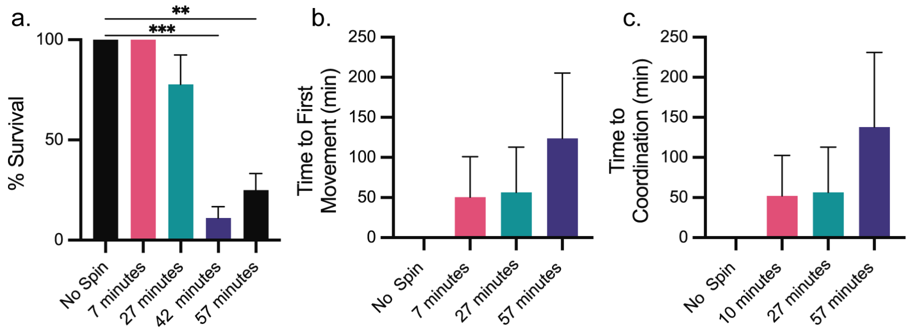


**Figure S12: 675,000 x g survival and recovery time courses.** a. Percent survival rate of tardigrades exposed to 675,000 x g for 7, 27, 42, and 57 minutes at maximum g-force. Two-way ANOVA column effects Dunnett's multiple comparisons test ***, p= 0.0005, **, p= -0.0036. b) Time to first movement reported as a function of HG exposure duration. c) Time to coordinated movement as a function of HG exposure duration. Kruskal-Wallis with Dunn's multiple comparisons test revealed no significant differences ~~from~~ between panels b and c.

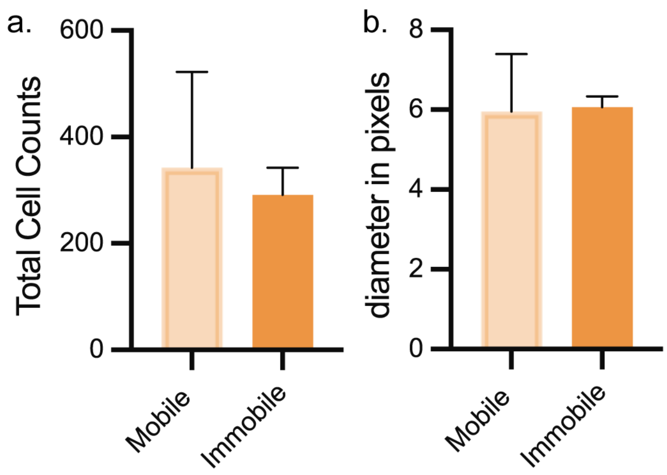


**Figure S13. Storage cell size and number post centrifugation.** a. Total Cell Counts measured obtained from mobile and immobile images. Welch's t-test shows no significant difference. b. Cell diameter of counted storage cells, Welch's t-test shows no significant difference.

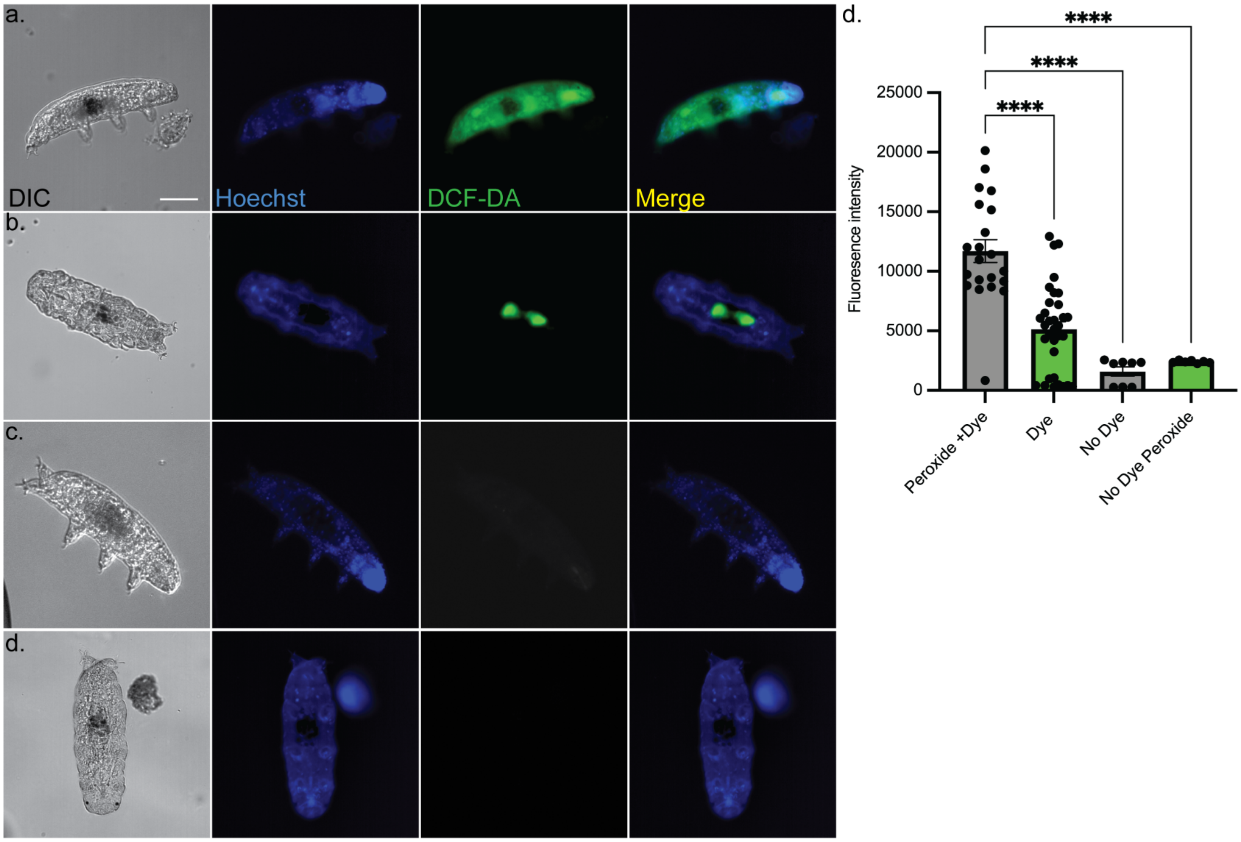


**Figure S14: ROS Assay enters cells and reliably reports peroxide activity in tardigrades.** a) DIC, Hoechst nuclear counterstain, DCF-DA, and fluorescence merged micrograph of Tardigrade treated with DFCDA and Hoechst prior to a 60-minute treatment with 3%peroxide. b) DIC, Hoechst, DCF-DA, and fluorescence merged image of Tardigrade treated with DCF-DA and Hoechst but not exposed to peroxide. c) Image panel suite of DIC, Hoechst, and DCF-DA of an animal exposed to peroxide in the absence of DCF-DA staining. d) Image panel of DIC, Hoechst, DFCDA, and merged image of animal exposed only to Hoechst, all scale bars are 50 μm.

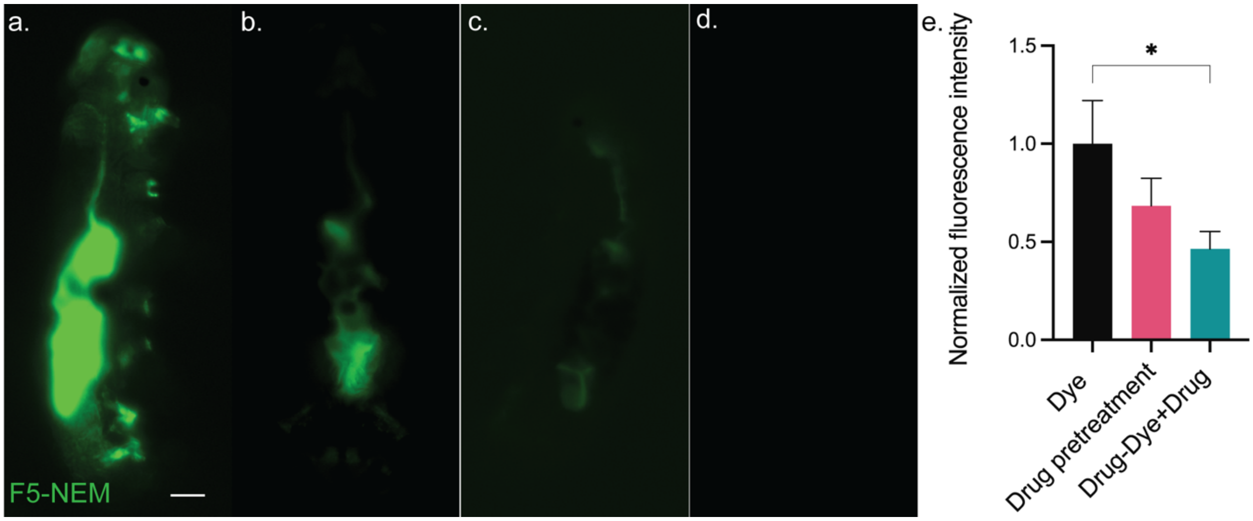


**Figure S15: NEM blocks available thiols, reducing NEM-dye conjugate labeling.** a. Fluorescence micrograph depicting tardigrades treated with F5-NEM, without NEM pretreatment. b. Epifluorescence image of a tardigrade pretreated with NEM, prior to exposure to F5-NEM. c. Epifluorescence image of an animal pretreated with NEM in the absence of F5-NEM. D. Fluorescence intensity of F5-NEM under dye only (black), NEM pretreatment (pink), and NEM pretreatment followed by equimolar presence of F5-NEM and NEM (green). Data represent average fluorescence intensity and S.E.M. across at least 13 animals per condition. Kruskal-Wallis test with Dunn's multiple comparison test, *, p=0.0373.

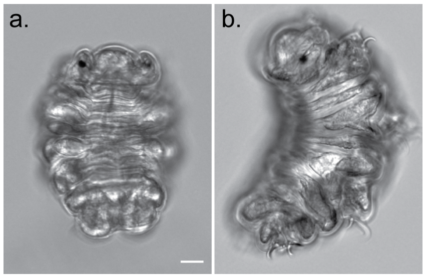


**Figure S16: Compete tun formation is inhibited by GSH pretreatment.** a. DIC Micrograph of tardigrade tun formation under control conditions, 0.5 M mannitol for 2 hours. Scale bar 10 µm. b. DIC micrograph of tardigrade pretreated with 30 mM GSH prior to hyperosmotic stress (0.5M mannitol for 2 hours.)

### Tables

**Table 1: A brief representation of maximum g-force resilience across the tree of life.**

| Species/Subject | Kingdom/Phylum | Max G-Force | Exposure Duration | Citation |
|---|---|---|---|---|
| *Paracoccus denitrificans* (bacteria) | Bacterium/Pseudomonadota | 403,627 g | 48 hours | (10) |
| *Panagrolaimus superbus & Caenorhabditis. elegans (nematodes)* | Animalia/Nematoda | 400,000 g | 1 hour | (11) |
| *Hypsibius exemplaris* (tardigrade) | Animalia/Tardigrada | 16,060 g | 1 minute | (12) |
| *Pisum sativum* (Pea, seed) | Plantae/Tracheophyta | 10,050 g | 48 hours | (13) |
| *Drosophila melanogaster* (Fruit fly) | Animalia/Arthropoda | Up to 9,000 g | 3 minutes | (14) |
| *Mus musculus* | Animalia/Chordata | 20 g | 10 minutes | (15) |
| *Homo sapiens* (with G-suit and AGSM*) | Animalia/Chordata | Up to 9 g | A few seconds | (16) |
| Pituophis melanoleucus (Snake) | Animalia/Chordata | 3 x g | 1 hour | (17) |

* AGSM: anti-G straining maneuvers

**Table 2: Stratification of cellular components invivo and invitro is dependent on density and size. Adapted from (18).**

| Cell Component | Component Density | Stratifying g-force in vivo | Duration | Citation |
|---|---|---|---|---|
| Nucleus, Glycogen | 0.8 $g/cm^3$ and 1.5 $g/cm^3$ | 19,000 g | 2 hours | Mateyko & Kopac 1954a, b, 1955, 1964) |
| Mitochondria, | 1.36 $g/cm^3$ | 19,000 g | 2 hours | Mateyko & Kopac 1954a, b, 1955, 1964) |
| Lysosomes, Peroxisomes and Cytoplasmic granules | 1.21 $g/cm^3$ , 1.15 $g/cm^3$, and 1.28 $g/cm^3$ | 19,000 g | 2 hours | Mateyko & Kopac 1954a, b, 1955, 1964) |
| ER and Microsome, | 1.16 $g/cm^3$ and 1.25 $g/cm^3$ | 110,000 g | 4 hours | Mateyko & Kopac (1965) |
| Ribosomes | 1.64 $g/cm^3$ | 95,000 g | 3 hours | David *et al.* (1961) |

**Table 3: Stratification within organellar compartments of animal cells**

| Within Organelle | | | | | |
|---|---|---|---|---|---|
| **Cell Component** | **Component Density** | **Stratifying g-force in vivo** | **Exposure Duration** | **Outcome** | **Citation** |
| Nuclei and Nucleous | 0.8 g/cm$^3$ and 2 g/cm$^3$ | 250,000 g | 10 min | Loss of nuclear envelope integrity | King & Beams (1934) |
| Golgi fractionation | 1.15 g/cm$^3$ | 200,000 g | 20 min | Separation into density dependent layers | Gatenby & Moussa (1949) |
| Mitochondria | 1.36 g/cm$^3$ | 400,000 g | 20 min | Separation into density dependent layers | Beams *et al.* (1960) |
